# Cholera toxin remodels intestinal immunity and suppresses systemic metabolism

**DOI:** 10.64898/2026.09.29.755509

**Authors:** Ana C. Roginski, Mahdieh Godazgar, Caio L. Salgado, Gourab Lahiri, Cheyanne Woodrow, Thomas H. McDermott, Bruna G. Costa Lima, Gina P. Rodriguez-Castano, Lee E. Voth-Gaeddert, Henrique Borges da Silva, Marcelo O. Dietrich, Miyeko D. Mana, Esther B. Florsheim

## Abstract

Cholera toxin (CTx) is a potent enterotoxin and widely used mucosal adjuvant, yet the acute host response to CTx itself, in the absence of a co-administered antigen, remains vague. In this study, we integrated physiological and metabolic phenotyping with intestinal transcriptomics, epithelial imaging, immune profiling, hepatic gene-expression analysis, and paired microbiome profiling to characterize the response to a single oral CTx exposure in mice. Within 24 h, CTx remodeled immune-cell composition in both epithelial and lamina propria compartments, with responses shaped by sex and genetic background. Recurrent changes included increased epithelial group 3 innate lymphoid cells, reduced lamina propria RORγt+ regulatory T cells, and decreased epithelial ILC1/NK cells in males of both strains. CTx also increased villus DCLK1+ tuft cells and reduced MUC2 immunoreactivity. Jejunal transcriptomics revealed induction of inflammatory defense, glycolytic, autophagy, and lipid-utilization programs together with suppression of proliferative and mitochondrial programs, consistent with coordinated stress adaptation. These local changes were accompanied by rapid restructuring of the fecal microbial community and by systemic hypometabolism, despite markedly different hypothermic responses across strains. CTx also elicited a selective hepatic stress and metabolic program, including conserved suppression of *Cyp7a1* across sexes and strains. Together, these findings show that CTx alone rapidly reorganizes the intestinal immune, epithelial, metabolic, and microbial environment while suppressing whole-body metabolism. This work provides an integrated framework for understanding the acute host response to CTx and for interpreting its widespread use as a mucosal adjuvant.

## INTRODUCTION

Ingested toxins disrupt intestinal homeostasis and elicit coordinated responses that can limit exposure or mitigate tissue damage, including fluid secretion, neuronal reflexes, metabolic adaptation, and detoxification^1,2^. These responses occur together with changes in epithelial and immune function, yet how an intestinal toxin reorganizes the local mucosal environment in the context of whole-body physiology remains incompletely understood. Defining these early responses is important for understanding how the host adapts to toxin-induced perturbation before exacerbated tissue injury or systemic disease develops.

Cholera toxin (CTx), an AB5 enterotoxin produced by *Vibrio cholerae*, provides a well-characterized model of intestinal intoxication. Binding of its pentameric B subunit to ganglioside GM1 facilitates delivery of the catalytic A1 subunit into the cytosol, where ADP-ribosylation of the stimulatory G-protein α subunit drives sustained cAMP signaling and CFTR-dependent chloride and fluid secretion^3–5^. Although these secretory effects are well established, their accompanying changes in intestinal tissue organization, immunity, metabolism, and host physiology are less clearly defined.

CTx is also one of the most widely used experimental mucosal adjuvants and has broad effects on innate and adaptive immunity. It can induce cytokine production, enhance antigen presentation, and promote dendritic cell maturation and migration to draining lymph nodes^6–10^. These activities support systemic and mucosal antibody responses and, depending on the experimental context, Th2-, Tr1-, or Th17-associated immunity^8,11–14^. Repeated oral administration of CTx with dietary proteins is therefore widely used to overcome oral tolerance and induce allergic sensitization in mice^15^. In this setting, IL-33 signaling through its receptor ST2 is required for robust allergic sensitization to peanut, implicating this pathway in CTx-induced mucosal type 2 immunity^16^. However, much of our understanding of CTx immunomodulation derives from *in vitro* studies or models in which CTx is administered together with food antigens. The acute response to CTx itself—before the development of antigen-specific adaptive immunity—remains poorly defined.

Here, we examined the early host response to a single oral exposure to CTx in C57BL/6 and BALB/c mice. We integrated physiological and metabolic measurements with hepatic gene-expression analysis, intestinal transcriptomics, epithelial imaging, immune profiling, and analysis of the fecal microbiota. We show that CTx induces systemic hypometabolism together with selective hepatic metabolic and stress responses, while rapidly remodeling the intestinal epithelial, immune, transcriptional, and microbial environment. These responses vary with sex and genetic background, revealing that the acute effects of CTx extend beyond fluid secretion. Together, our findings provide an integrated view of the early host response to a widely used mucosal toxin and adjuvant.

## RESULTS

### Acute oral exposure to cholera toxin alters physiology and suppresses whole-body metabolism

Cholera toxin (CTx) is a potent enterotoxin that induces rapid fluid secretion, and while its cellular responses are well-characterized, the early tissue-level responses to CTx exposure remain incompletely defined. We established an acute oral exposure model in mice to capture the physiological and intestinal responses to this clinically relevant toxin. C57BL/6 mice of both sexes received a single nonlethal intragastric dose of 0.8 or 1.6 mg/kg CTx during their active phase (dark cycle, ZT14) and were monitored for physiological symptoms for up to 6 h (Figure 1a). The 6 h endpoint was chosen to capture the initiation of the secretory response, after mucosal cAMP is elevated and secondary transcriptional programs are engaged^17^. We selected the 0.8 mg/kg dose based on previous studies reporting CTx-induced diarrhea and mild intestinal fluid secretion^18^. Moreover, we tested the higher dose of CTx (1.6 mg/kg) to determine whether more severe toxin exposure would elicit distinct physiological and intestinal responses.

**Figure 1.**
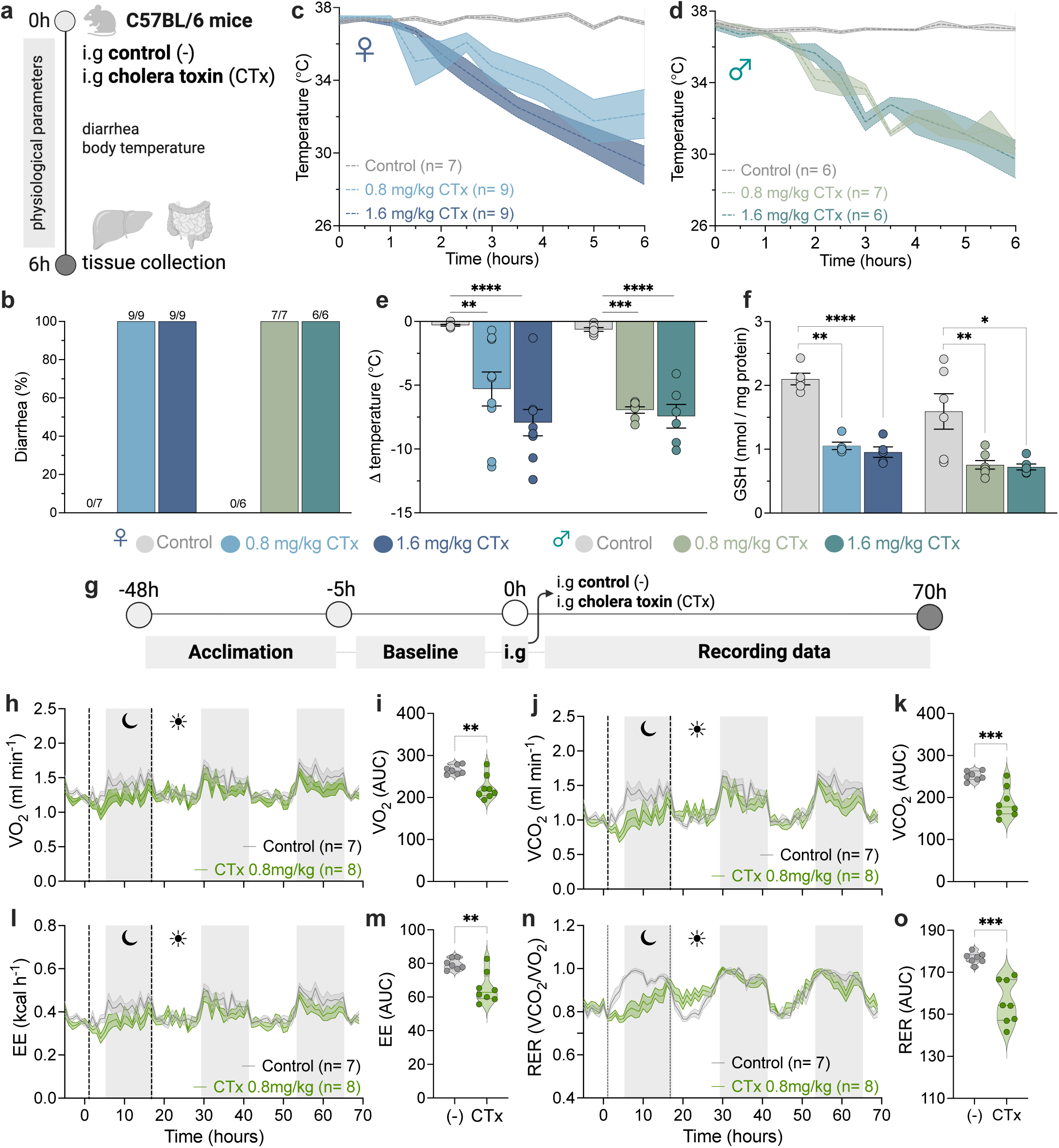
Acute oral exposure to CTx induces hypothermia, intestinal fuild accumulation and systemic metabolic reprogramming in C57BL/6 mice. C57BL/6 mice (6-8 weeks old, both sexes) were used to assess physiological responses to oral cholera toxin (CTx). **(a)** Schematic of the experimental protocol for acute CTx exposure and assessment of physiological responses. **(b)** Diarrhea occurrence; blue bars represent females, green bars represent males, and gray bars represent sex-matched controls (n= 6-9 mice per group). **(c-d)** Rectal temperature over time following oral CTx exposure; **(c)** females and **(d)** males. **(e)** Maximum decrease in rectal temperature; blue bars represent females, green bars represent males and gray bars represent sex-matched controls (n= 6-9 mice per group). **(f)** Reduced glutathione (GSH) levels in the jejunum; blue bars represent females, green bars represent males, and gray bars represent sex-matched controls (n= 6-9 mice per group). For metabolic evaluations, C57BL/6 male mice were used (n= 7-8 mice per group). **(g)** Schematic of the experimental protocol for metabolic assessment following oral CTx exposure. **(h)** Oxygen consumption over time and **(i)** area under the curve (AUC) from 1 to 16.6 h post-exposure. **(j)** Carbon dioxide production over time and **(k)** corresponding AUC from 1 to 16.6 h post-exposure. **(l)** Energy expenditure (EE) over time and **(m)** corresponding AUC from 1 to 16.6 h post-exposure. **(n)** Respiratory Exchange Ratio (RER) over time and **(o)** corresponding AUC from 1 to 16.6 h post-exposure. Data are presented as mean ± s.e.m. Statistical significance was determined using unpaired two-tailed *Student’s t*-test comparing control to CTx-treated group. * p ≤ 0.05, ** p ≤ 0.01, *** p ≤ 0.001, **** p ≤ 0.0001. Each panel represents data from at least two independent experiments. In panels **h**, **j**, **l**, **n**, gray shaded areas indicate the dark cycle, and white areas indicatethe light cycle; dashed lines denote AUC interval. Schematics in **a** and **g** were created with BioRender.com.

Following oral CTx administration, we observed watery diarrhea in all treated mice, as expected (Figure 1b). Interestingly, we observed that CTx induced marked hypothermia (Figures 1c-d). Body temperature dropped from baseline by an average of 5.3°C and 7.9°C in females exposed to 0.8 and 1.6 mg/kg of CTx, respectively. In males, the corresponding average decreases were 6.9°C and 7.4°C, respectively, compared to controls (Figure 1e). In both sexes, body temperature continued to decline throughout the 6 h monitoring period. CTx also reduced jejunal levels of glutathione (GSH) (Figure 1f), consistent with altered redox homeostasis, as GSH is one of the body’s primary endogenous antioxidants^19^. Finally, at 6 h post-exposure all CTx-treated mice appeared hypoactive and less responsive to handling than controls. These observations were qualitative and were not formally scored.

Because body temperature is closely linked to metabolic function, reductions in metabolic activity can be accompanied by decreased heat production and lower body temperature^20^. Conversely, decreases in body temperature can also suppress metabolic processes by slowing enzyme-dependent biochemical reactions, thereby reinforcing a hypometabolic, energy-conserving state^21^. To test if CTx induces metabolic alterations, we used a high-resolution metabolic phenotyping system (Promethion) and evaluated individually caged mice. C57BL/6 males were chosen because they exhibited a stronger hypothermic response to the lower CTx dose. Mice were acclimated to the cages for 48 h, and we started recording baseline metabolic parameters 5 h before CTx exposure. After oral toxin administration, mice were monitored for oxygen consumption (VO_2_), carbon dioxide production (VCO_2_), total energy expenditure (EE), and respiratory exchange ratio (RER) for up to 70 h (Figure 1g). Overall, CTx induced a global reduction in metabolic activity, with decreases of 15.8% in the VO_2_ (Figures 1h-i), 25.1% in the VCO_2_ (Figures 1j-k), 17.8% in EE (Figures 1l-m), and 11.8% in RER (Figures 1n-o). The hypometabolism started approximately 1 h after the exposure and lasted for up to 16.6 h (highlighted by dashed lines in Figures 1h, j, l, n). In control mice, the RER values show the daily cycle of fuel utilization, with predominant fat oxidation (RER≈0.8) during the light phase (inactive period in rodents, white areas), followed by a shift toward carbohydrate oxidation (RER≈1.0) during the dark phase (active period, gray shaded areas) (Figure 1n). However, CTx-treated mice delayed this shift from fat to carbohydrate utilization for the first 30 h post-exposure, approximately, after which treated mice resumed a pattern similar to controls (Figure 1n).

To evaluate whether CTx effects are strain-specific, we determined the physiological and metabolic consequences of oral CTx exposure in BALB/c mice (Figure 2a). In this strain, CTx did not induce diarrhea during the 6 h of monitoring, although intestinal fluid accumulation occurred in both the small intestine and cecum of all treated mice (Figure 2b). In addition, CTx caused only minimal hypothermia in females when exposed to 1.6 mg/kg of toxin (a drop of −0.94°C; Figures 2c, e), and a slightly greater drop in males (−1.1°C and −1.7°C, at the lower and higher CTx dose, respectively; Figures 2d, e). Notably, CTx reduced jejunal GSH levels in females, but not in males (Figure 2f). BALB/c mice exhibited less pronounced hypothermia and no observed diarrhea during the initial 6-h observation period.

**Figure 2.**
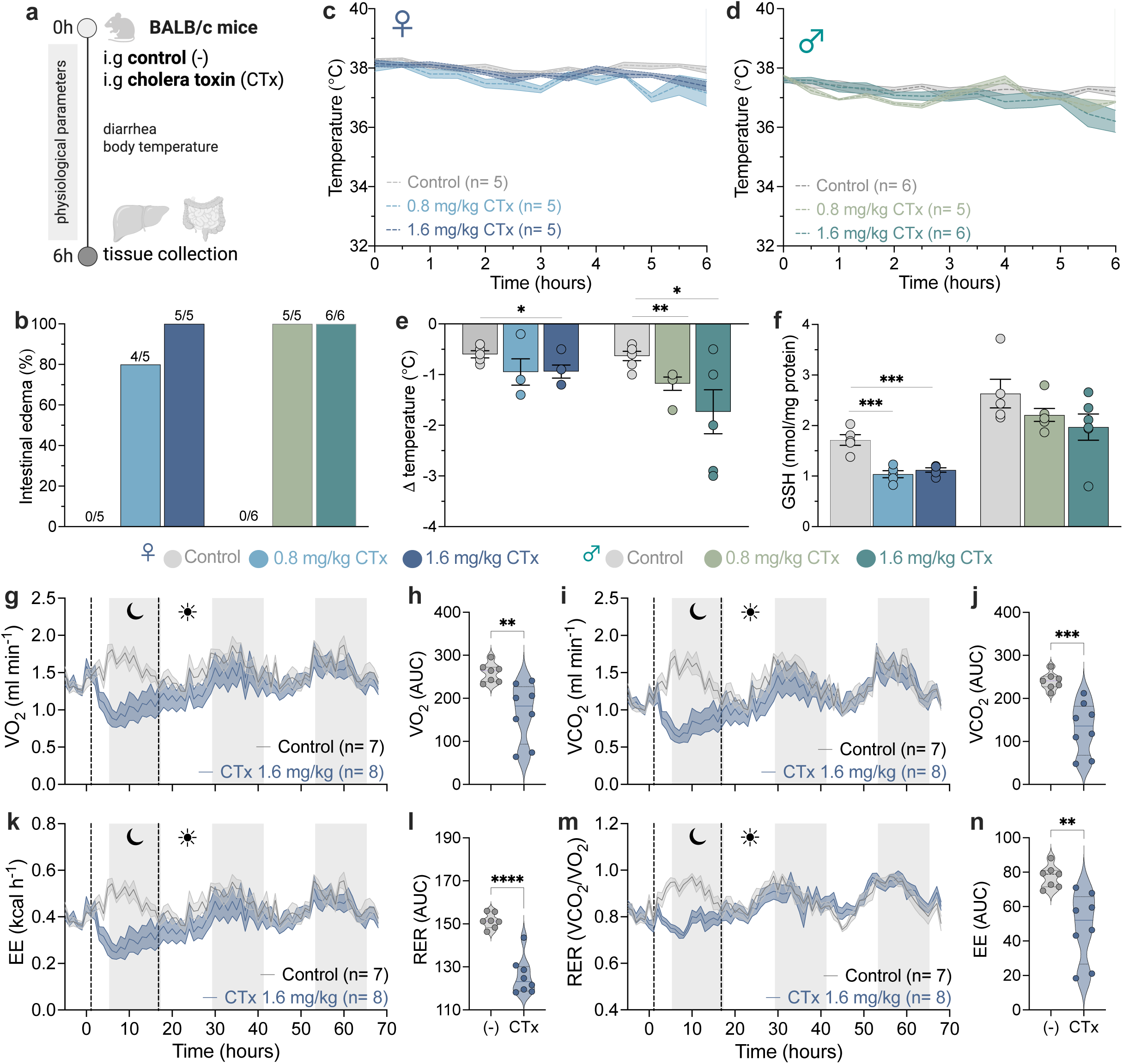
Acute oral exposure to CTx triggers intestinal fluid accumulation and systemic metabolic adaptations in BALB/c mice. BALB/c mice (6-8 weeks old, both sexes) were used to evaluate physiological responses to oral cholera toxin (CTx). **(a)** Schematic of the experimental protocol. **(b)** Quantification of small intestinal fluid accumulation; blue bars indicate females, green bars indicate males, and gray bars indicate sex-matched controls (n= 5-6 mice per group). **(c-d)** Rectal temperature over time following oral CTx exposure; **(c)** females and **(d)** males. **(e)** Maximum decrease in rectal temperature; blue bars represent females, green bars represent males, and gray bars represent sex-matched controls (n= 5-6 mice per group). **(f)** Reduced glutathione (GSH) levels in the jejunum; blue bars represent females, green bars represent males, and gray bars represent sex-matched controls (n= 5-6 mice per group). For metabolic evaluations, BALB/c female mice were used (n= 7-8 mice per group). **(g)** Oxygen consumption over time and **(h)** area under the curve (AUC) from 3 to 16.6 h post-exposure. **(i)** Carbon dioxide production over time and **(j)** corresponding AUC from 3 to 16.6 h post-exposure. **(k)** Energy Expenditure (EE) over time and **(l)** corresponding AUC from 3 to 16.6 h post-exposure. **(m)** Respiratory Exchange Ratio (RER) over time and **(n)** corresponding AUC from 3 to 16.6 h post-exposure. Data are presented as mean ± s.e.m. Statistical significance was determined using unpaired two-tailed *Student’s t*-test comparing control to CTx-treated group. * p ≤ 0.05, ** p ≤ 0.01, *** p ≤ 0.001, **** p ≤ 0.0001. Each panel represents data from at least two independent experiments. In panels **g**, **i**, **k**, **m**, gray shaded areas indicate the dark cycle, and white areas indicate the light cycle; dashed lines denote AUC interval.

We next hypothesized that the strain-dependent differences observed in physiological outcomes (e.g., hypothermia and diarrhea) could be associated with variation in the expression of the intestinal GM1 ganglioside receptor, required for CTx internalization and pathogenesis^5^. To test this, we quantified the intestinal expression of three key genes encoding enzymes involved in the GM1 biosynthetic pathway (*St3gal5*, *B4galnt1*, *B3galt4*) and one gene encoding an enzyme responsible for GM1 degradation (*Glb1*) in jejunal samples using quantitative real-time PCR (qPCR) (Suppl. Figure 1a). Two-way ANOVA revealed no statistically significant differences between males and females within either strain for any of the genes analyzed, and therefore, data from both sexes were combined for strain comparisons. BALB/c mice exhibited significantly higher jejunal expression of *St3gal5* (GM3 synthase) and *B4galnt1* (GM2/GD2 synthase), encoding the enzymes responsible for the sequential conversion of lactosylceramide to GM3 and GM3 to GM2, respectively (Suppl. Figure 1b-c). In contrast, the expression of *B3galt4* (GM1 synthase), which catalyzes the terminal biosynthetic step converting GM2 into GM1, was significantly lower in BALB/c mice compared to C57BL/6 mice (Suppl. Figure 1d). No strain- or sex-related differences were observed in the expression of *Glb1* (β-galactosidase), the enzyme responsible for lysosomal GM1 degradation (Suppl. Figure 1e). Taken together, these findings suggest that, despite an apparent upregulation of the upstream ganglioside biosynthetic pathway, BALB/c mice may exhibit reduced intestinal GM1 synthesis relative to C57BL/6 mice. These findings identify strain-associated differences in the expression of genes involved in ganglioside metabolism; their consequences for intestinal GM1 abundance and toxin uptake remain unresolved.

Analysis of the metabolic responses in BALB/c females revealed that CTx also suppressed metabolic activity with reductions of 35.6% in VO_2_ (Figures 2g-h), 35.6% in VCO_2_ (Figures 2i-j), 37.6% in EE (Figures 2k-l), and 16.9% in RER (Figures 2m-n). Similarly, CTx disrupted the normal light-to-dark transition in fuel utilization, delaying the shift from fat to carbohydrate oxidation for up to ∼20 h post-exposure (Figure 2m).

Together, these findings show that acute oral CTx exposure alters intestinal physiology and suppresses whole-body metabolism. Metabolic suppression was observed in both C57BL/6 males and BALB/c females, despite markedly different hypothermic responses in the physiological cohorts. These observations suggest that pronounced hypothermia is not required for the metabolic phenotype, but the contributions of strain, sex, exposure conditions, and cohort differences cannot be separated by these comparisons.

### Cholera toxin induces selective hepatic stress and metabolic transcriptional responses

Physiological responses to harmful exposures extend beyond the neutralization or elimination of the noxious agent. Changes in energy expenditure, substrate utilization, and tissue metabolism can also influence the capacity to overcome physiological stress^22,23^. The liver is central to these responses via fuel metabolism, xenobiotic processing, and the production of inflammatory and endocrine mediators^24,25^. The acute phase responses induced by systemic exposure to lipopolysaccharide (LPS) in the liver, for example, is a hallmark of inflammation, yet we know little about the effects of ingested toxins in promoting these acute phase reactions. We next assessed hepatic gene expression 6 h after oral CTx exposure, focusing on markers of acute-phase response, oxidative stress, xenobiotic metabolism, and inflammation.

In C57BL/6 males, oral CTx reduced hepatic C-reactive protein (*Crp*) expression, whereas no significant change in serum amyloid A (*Saa1*) was detected (Figure 3a and Suppl Figure 2a). Interleukin-1 β (*Il1b*) and interleukin-6 (*Il6*) expression did not differ significantly between groups, while tumor necrosis factor α (*Tnf*) expression decreased (Suppl Figure 2b-c and Figure 3b). Thus, the hepatic transcripts examined did not show coordinated induction of a canonical acute-phase-associated response at this time point.

**Figure 3.**
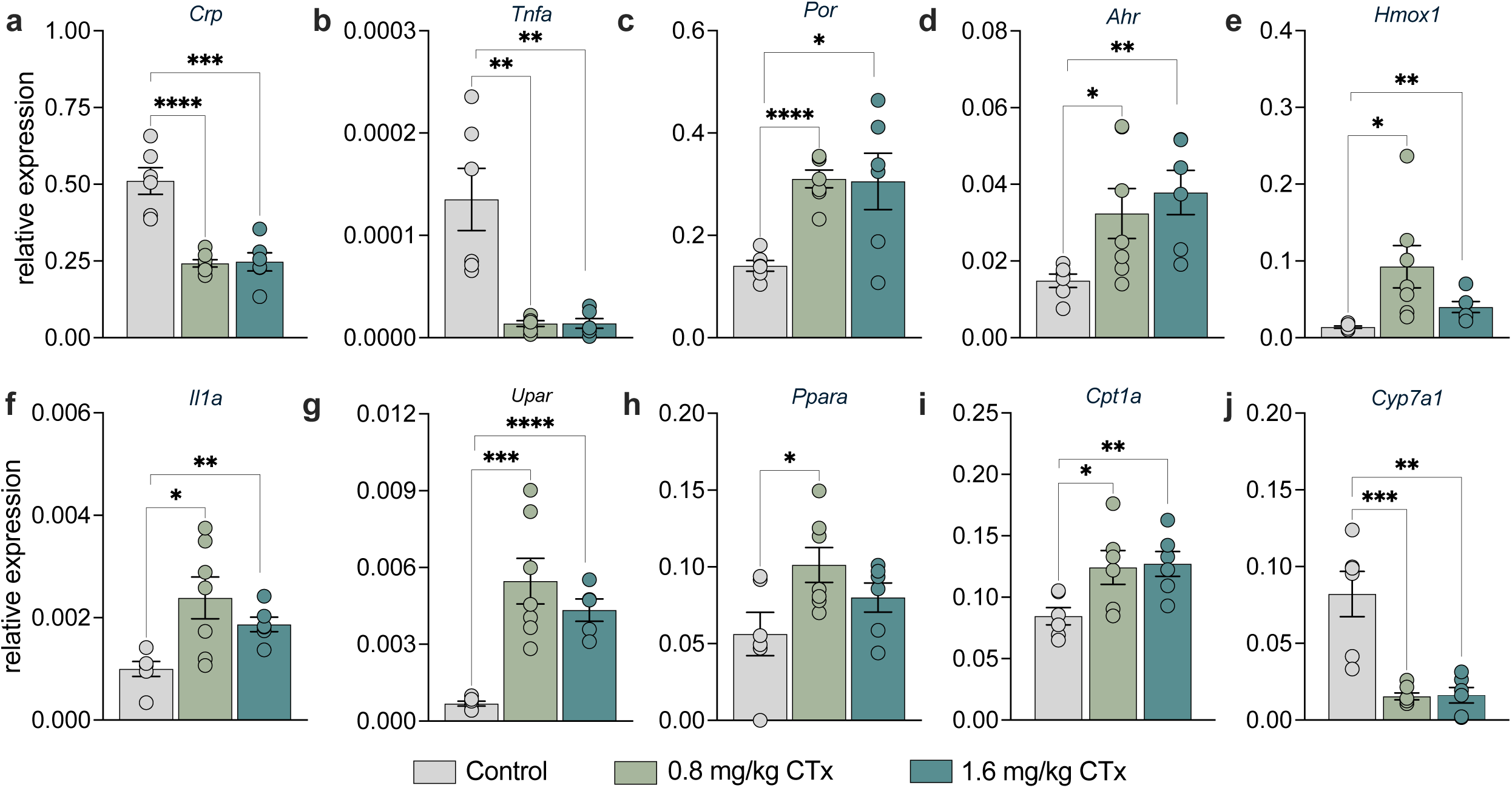
Oral CTx induces early hepatic detoxification and stress-responses gene expression. Livers from C57BL/6 male mice were collected 6 h after a single oral exposure to cholera toxin (CTx) and analyzed by quantitative real-time PCR (n= 6-7 mice per group). CTx significantly altered hepatic expression of: **(a)** C-reactive protein (*Crp*), **(b)** cytochrome P450 oxidoreductase (*Por*), **(c)** aryl hydrocarbon receptor (*Ahr*), **(d)** urokinase plasminogen activator surface receptor (*Upar*), **(e)** heme oxygenase 1 (*Hmox1*), **(f)** interleukin-1α (*Il1a*), **(g)** tumor necrosis factor α (*Tnfa*), **(h)** peroxisome proliferator-activated receptor α (*Ppara*), **(i)** carnitine palmitoyltransferase 1A (*Cpt1a*), and **(j)** cholesterol 7α-hydroxylase (*Cyp7a1*). mRNA levels were normalized to the housekeeping gene *Rpl13a* and are presented as relative expression (mean ± s.e.m). Statistical significance was assessed using unpaired two-tailed *Student’s t*-test comparing control to CTx-treated group. * p ≤ 0.05, ** p ≤ 0.01, *** p ≤ 0.001, **** p ≤ 0.0001. Each panel represents pooled data from three independent experiments.

Genes associated with xenobiotic and stress responses were differentially regulated by CTx. Cytochrome P450 oxidoreductase (*Por)* (Figure 3c) and the aryl hydrocarbon receptor (*Ahr*) (Figure 3d) were upregulated. However, this increase in *Ahr* expression was not accompanied by changes in its canonical downstream targets, AHR repressor (*Ahrr*) or cytochrome P450 1A1 (*Cyp1a1*) (Suppl. Figure 2d-e), indicating increased *Ahr* transcript abundance without coordinated induction of the measured canonical targets. Consistent with a stress-related response, the oxidative stress marker heme oxygenase 1 (*Hmox1*) was upregulated (Figure 3e). In contrast, no significant change was detected for other antioxidant-related genes, including 5-oxoprolinase (*Oplah*), glutathione-disulfide reductase (*Gsr*), superoxide dismutase 1 (*Sod1*), and glutathione peroxidase 1 (*Gpx1*) (Suppl. Figure 2f-i).

CTx increased expression of IL-1α (*Il1a*) (Figure 3f) and urokinase plasminogen activator surface receptor (*Plaur, or uPar*) (Figure 3g) and of genes associated with fatty acid oxidation, including peroxisome proliferator-activated receptor α (*Ppara*), and carnitine palmitoyltransferase 1A (*Cpt1a*) (Figure 3h-i). Conversely, CTx downregulated the cholesterol 7α-hydroxylase (*Cyp7a1*) (Figure 3j), the rate-limiting enzyme in bile acid synthesis. Notably, suppression of *Cyp7a1* has been shown during sustained inflammation^26^. No significant changes in liver-derived fibroblast growth factor 21, *Fgf2*1, or *Klb*, which encodes the FGF21 coreceptor β-klotho, were detected (Suppl. Figure 2j-k). Interestingly, FGF21 was previously reported to maintain thermoregulation during bacterial inflammation^27^.

Together, these findings indicate that acute CTx exposure in C57BL/6 males elicits a selective hepatic response characterized by the engagement of stress and metabolic pathways alongside limited activation of canonical inflammatory signaling. We next asked whether this response is conserved across sexes and genetic backgrounds, or whether these variables shape the early systemic response to CTx.

In C57BL/6 females, CTx elicited a partially overlapping hepatic response compared to males, with *Upar* and *Por* upregulation and *Crp* and *Cyp7a1* downregulation. However, several stress- and inflammation-associated genes that were induced or unchanged in males, including *Hmox1*, *Oplah*, *Gsr*, *Gpx1*, and *Il1b,* were downregulated in females. Unlike the response observed in males, hepatic *Fgf21* expression increased in C57BL/6 females, whereas *Klb* remained unchanged. Finally, other genes including *Ppara, Il6*, *Tnfa*, *Ahr*, *Sod1*, *Il1a*, and *Cpt1a* remained unchanged (Suppl. Figure 3).

In BALB/c mice, CTx consistently induced *Upar* and *Il1b* and suppressed *Cyp7a1*, across sexes. Hepatic *Fgf2*1 expression also increased in both males and females, whereas *Klb* remained unchanged. Additional transcriptional changes were limited and sex-dependent, with females showing downregulation of *Ahr* and *Por*, and males exhibiting upregulation of *Il1a*, while most other genes remained unchanged (Suppl. Figure 4).

Collectively, these findings identify a selective hepatic transcriptional response to acute oral CTx exposure. *uPar* induction and *Cyp7a1* suppression were detected in each sex–strain group examined, whereas the patterns of detected changes in stress-associated genes and metabolic regulators differed across groups.

### Cholera toxin alters intestinal stress-response and metabolic gene expression

The primary site of action of CTx is the small intestine. To characterize the early intestinal response to toxin exposure, we performed bulk RNA sequencing of proximal jejunum samples collected 6 hours after CTx administration in male C57BL/6 mice. Principal component analysis (PCA) revealed a clear separation between control and CTx-treated samples, indicating a robust and coordinated transcriptional response to the toxin (Figure 4a). Consistent with this separation, differential expression analysis identified 2,177 significantly differentially expressed genes (DEGs), with 1,065 upregulated and 1,112 downregulated (Figure 4b). Among the 10 most differentially expressed transcripts of the small intestine, *Mt1*, *Mt2*, *Ddit4*, *Fut4*, *Akr1b7*, *Cirbp*, *Pnpla2*, *Slc25a28*, and *Sesn1* were highly upregulated, while *Apol7c* was downregulated (Figure 4b). Several of these are linked to cellular stress responses, metabolic regulation, and epithelial defense mechanisms, suggesting that acute toxin exposure might rapidly alter pathways involved in intestinal tissue homeostasis.

**Figure 4.**
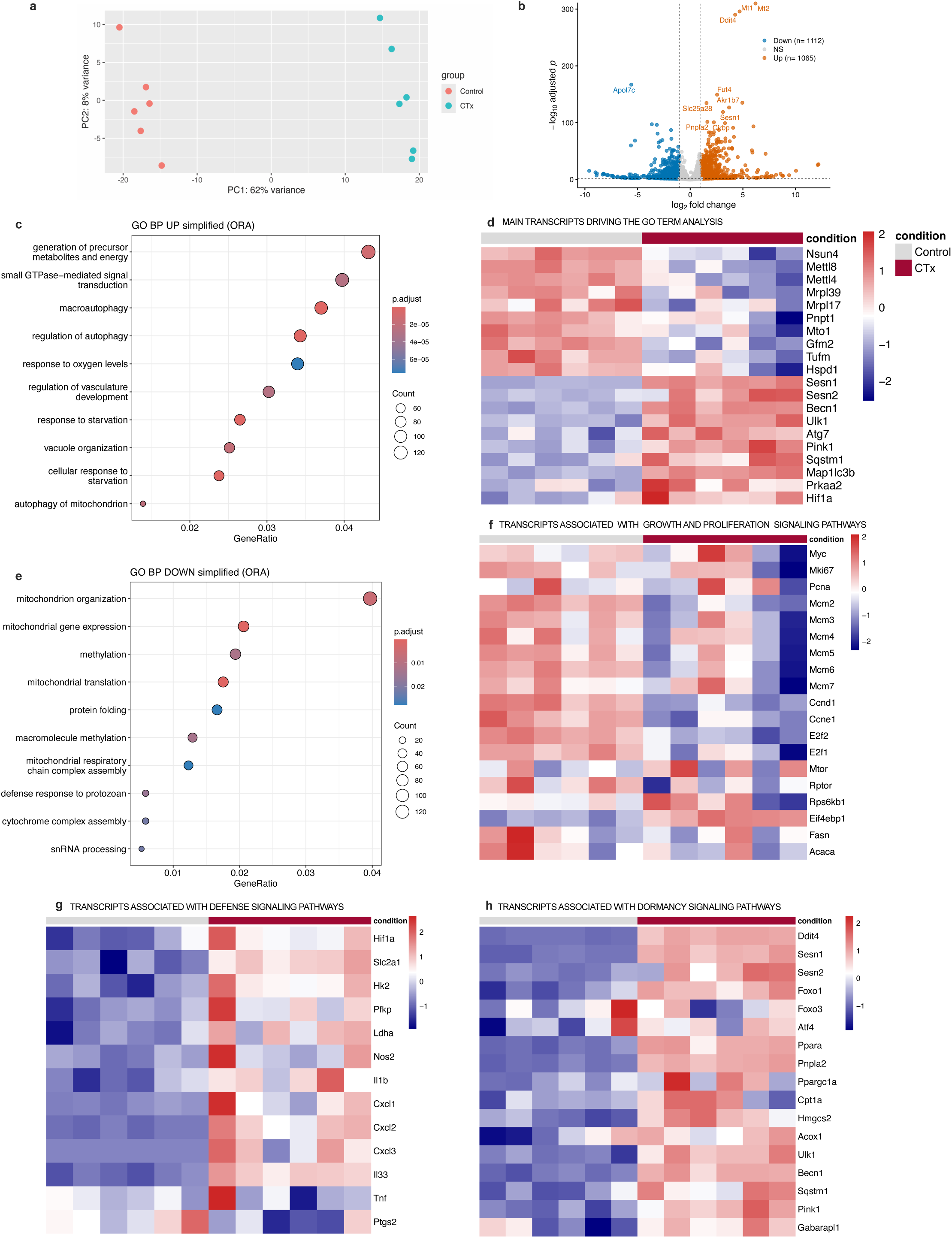
CTx exposure triggers intestinal defense and dormancy-like maintenance programs. Proximal jejunum tissue from C57BL/6 male mice was collected 6 h after a single oral exposure to cholera toxin (CTx) and analyzed by bulk RNA sequencing (n= 6 mice per group). A total of 2,177 transcripts were differentially expressed (adjusted *P* <0.05, |log2 fold change| ≥1), with 1,065 upregulated and 1,112 downregulated in CTx-treated mice compared to controls. **(a)** Principal component analysis (PCA) showing separation between control and CTx-treated samples. **(b)** Volcano plot of differentially expressed transcripts. **(c–e)** Gene Ontology (GO) enrichment analysis of differentially expressed transcripts (adjusted *P* < 0.05, |log2 fold change| = 0). **(c)** Top enriched biological processes among upregulated transcripts. **(d)** Heatmap showing the expression patterns of transcripts driving enriched GO biological processes. **(e)** Top enriched biological processes among downregulated transcripts. **(f-h)** Heatmap of transcripts associated with **(f)** growth and proliferation, **(g)** defense, and **(h)** dormancy-related signaling pathways identified by RNA sequencing. Data are pooled from at three independent experiments. Heatmaps in (**d, f-h**) show row-wise Z-score-normalized expression values.

Gene ontology (GO) enrichment analysis of differentially expressed transcripts revealed distinct biological programs associated with downregulated and upregulated transcripts following CTx exposure. Upregulated transcripts were significantly enriched in processes related to autophagy and vacuole organization, together with broader stress and metabolic responses, including cellular response to starvation, generation of precursor metabolites and energy, response to oxygen levels, and small GTPase–mediated signal transduction (Figure 4c). Enrichment was associated with genes including canonical regulators of autophagy and metabolic stress, including *Becn1*, *Ulk1*, *Atg7*, *Pink1*, *Sqstm1*, and the stress-response genes *Sesn1* and *Sesn2* (Figure 4d). In contrast, downregulated genes were strongly enriched in mitochondrial-related processes, as well as pathways related to protein folding, snRNA processing, and macromolecule methylation (Figure 4e). These pathways were driven by transcripts encoding mitochondrial ribosomal proteins and components of the mitochondrial translation machinery, such as *Mrpl* and *Mrps* family members, along with mitochondrial RNA-processing and methylation factors including *Nsun4*, *Mettl8*, and *Mettl4* (Figure 4d). Collectively, these expression changes identify a stress-associated transcriptional signature involving autophagy-related genes and reduced expression of genes associated with mitochondrial and biosynthetic processes.

We next interpreted the transcriptional changes within a framework of growth, defense, and maintenance programs^28^. Because the relevant signaling pathways are extensively regulated after transcription, we examined associated gene-expression signatures rather than pathway activity. CTx reduced the expression of several regulators of cell-cycle progression and DNA replication, including *Ccnd1*, *Ccne1*, *E2f1*, *E2f2*, *Mcm2*, *Mcm5*, and *Mcm6*, consistent with reduced investment in proliferative processes (Figure 4f). In parallel, CTx induced a defense-associated glycolytic program characterized by increased expression of *Hif1*a, *Slc2a1*, *Hk2*, *Pfkp*, and *Ldha*, accompanied by induction of inflammatory mediators including *Nos2*, *Il1b*, *Cxcl1-3,* and *Il33* (Figure 4g). Notably, CTx also induced a prominent stress- and nutrient-responsive transcriptional program associated with dormancy-like metabolic adaptation, including *Ddit4*, *Sesn1*, and *Sesn2*, but also *Foxo1*, and *Ppara*. This response was accompanied by increased expression of genes involved in lipid mobilization and utilization, such as *Pnpla2*, *Ppargc1a*, *Cpt1a*, and *Hmgcs2*, as well as transcripts associated with autophagy and mitochondrial quality control, including *Ulk1*, *Becn1*, *Sqstm1*, *Pink1*, and *Gabarapl1* (Figure 4h). These findings are consistent with reduced expression of proliferative programs alongside inflammatory, glycolytic, and maintenance-associated transcriptional responses.

### Cholera toxin alters intestinal epithelial cell markers

We next evaluated the effects of CTx on intestinal epithelial cell composition by immunofluorescence 24 hours after toxin exposure, in male C57BL/6 mice (Figure 5a). Immunostaining revealed a marked increase in the number of DCLK1^+^ cells, a marker of tuft cells, along the villus epithelium in CTx-treated animals compared to controls (Figure 5b), whereas no differences were observed in the crypts (Figure 5c). In contrast, fewer MUC2^+^ epithelial cells were detected (Figure 5d-e). Enteroendocrine (CHGA^+^ cells) and Paneth (LYZ1^+^ cells) cells per villus or crypt did not differ between groups (Figure 5f-h).

**Figure 5.**
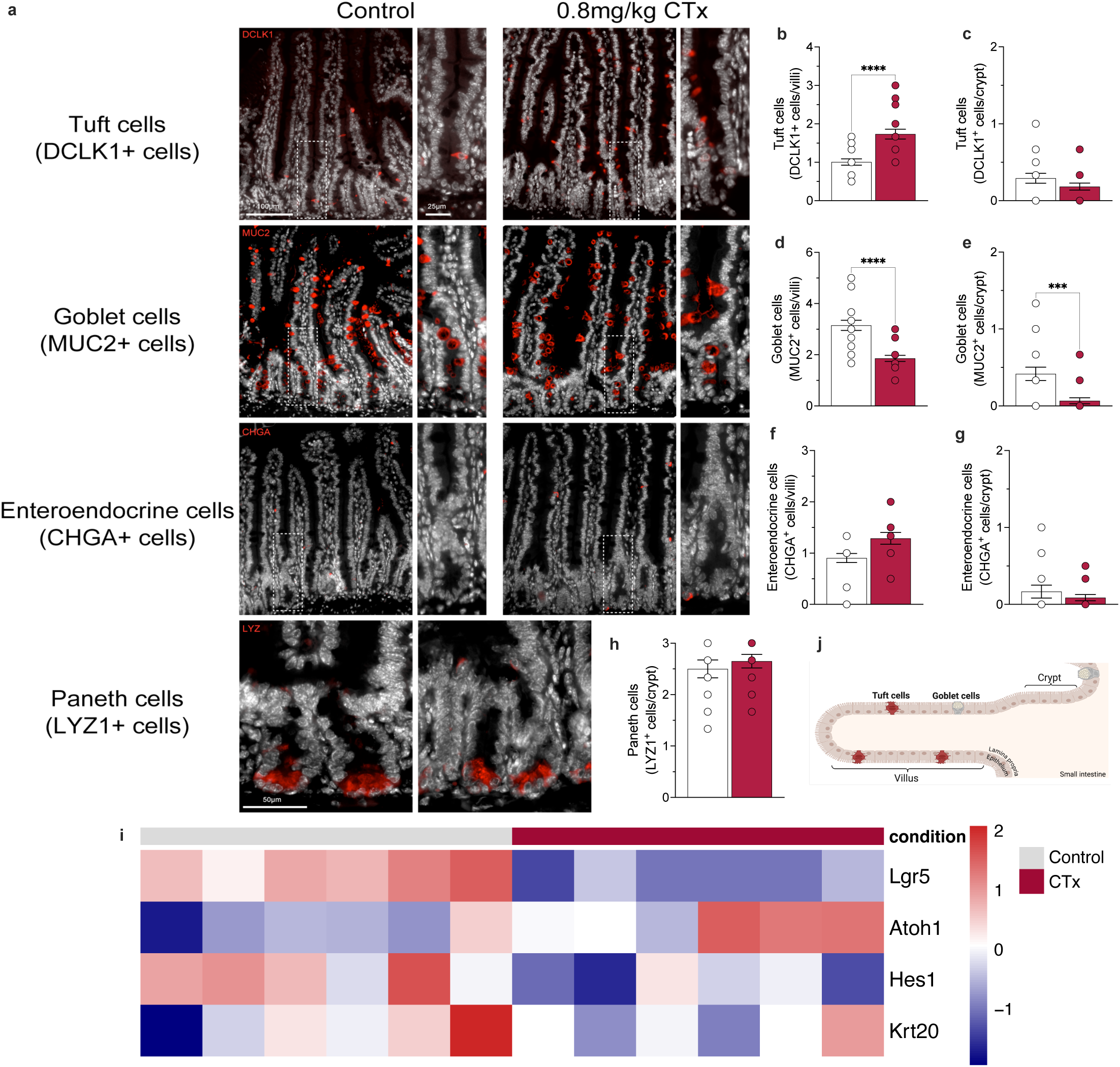
Cholera toxin alters intestinal epithelial cell composition. Jejunum samples from C57BL/6 male mice were analyzed. **(a)** Representative immunofluorescence images of jejunal villi and crypt regions collected 24 h after a single oral exposure to CTx. Tissues were fixed in 4% paraformaldehyde overnight, embedded in paraffin, and processed for immunofluorescence staining. **(a)** Sections were immunolabeled with antibodies against DCLK1 (tuft cell marker, red), MUC2 (goblet cell marker, red), CHGA (enteroendocrine cell marker, red), and LYZ1 (Paneth cell marker, red), counterstained with DAPI (white), and imaged using fluorescence microscopy at 20x magnification (n= 3 mice per group). **(b–h)** Quantification of epithelial cell populations in the small intestine 24 h after CTx exposure, showing tuft cell numbers **(b)** per villi and **(c)** per crypt, goblet cell numbers **(d)** per villi, and **(e)** per crypt, enteroendocrine cell numbers **(f)** per villi, and **(g)** per crypt, and paneth cell numbers **(h)** per crypt. Quantification was based on the number of marker-positive cells per 20 villi or crypts across multiple sections per animal (n= 3 mice per group). **(i)** RNA sequencing analysis of proximal jejunum collected 6 h after CTx exposure (n= 6 mice per group). Normalized expression levels (row-wise Z-score) of representative markers associated with epithelial stemness and differentiation markers are shown. **(j)** Summary schematic of immunofluorescence findings. Data are presented as mean ± s.e.m. Statistical significance was determined using unpaired two-tailed *Student’s t*-test comparing control and CTx-treated group. *** p≤ 0.001, **** p≤ 0.0001. <u>Abbreviations</u>: Lgr5, leucine rich repeat containing G protein-coupled receptor 5; Atoh1, atonal bHLH transcription factor 1; Hes1, hes family bHLH transcription factor 1; and Krt20, keratin 20.

To characterize early alterations in intestinal epithelial cell composition, we analyzed key markers associated with stem cells (*Lgr5*), secretory-lineage specification (*Atoh1*), absorptive-lineage specification (*Hes1*), and epithelial differentiation/maturation (*Krt20*), by RNA-sequencing 6 h post toxin administration. This analysis revealed decreased expression of *Lgr5* and *Hes1*, accompanied by increased *Atoh1* expression, whereas *Krt20* remained unchanged (Figure 5i). These changes are consistent with altered expression of genes associated with epithelial stem-cell and lineage regulation. Together with the later immunostaining findings, they indicate an epithelial response to CTx, although the relationship between the early transcriptional changes and the 24-h cellular phenotype remains unresolved.

### Cholera toxin induces an immune shift in the small intestine

CTx is a potent mucosal adjuvant widely used to induce adaptive immunity in food allergy models. Consequently, most of the knowledge regarding the immune effects of CTx derives from studies in which this toxin is co-administered with a food antigen. Although CTx is known to induce activation and trigger DC migration from the intestinal tissue to the regional draining lymph nodes^10,29^, its effects on other immune cell populations remain largely unknown. To address this, we evaluated immune cell frequencies of the epithelial and lamina propria compartments of the small intestine in male C57BL/6 mice 24 h after CTx exposure using full-spectrum flow cytometry.

In the epithelial compartment, CTx rapidly increased the frequencies of group 3 innate lymphoid cells (ILC3s), B cells, CD4⁺ T cells, monocytes, group 2 innate lymphoid cells (ILC2s), and Th17 cells. In contrast, the frequencies of γδ T cells, group 1 innate lymphoid cells (ILC1/NK population), and total T cells were reduced compared with control mice exposed to vehicle solution alone (Figure 6a-i). In the lamina propria, CTx similarly induced selective changes in immune populations, characterized by increased frequencies of CD8⁺ T cells, B cells, and CD103⁺CD11b⁺ DCs. Conversely, CD103⁺CD11b-DC, macrophages, Foxp3⁺ regulatory T cells, RORγT⁺ regulatory T cells, and total T cells were decreased (Figure 6j-q).

**Figure 6.**
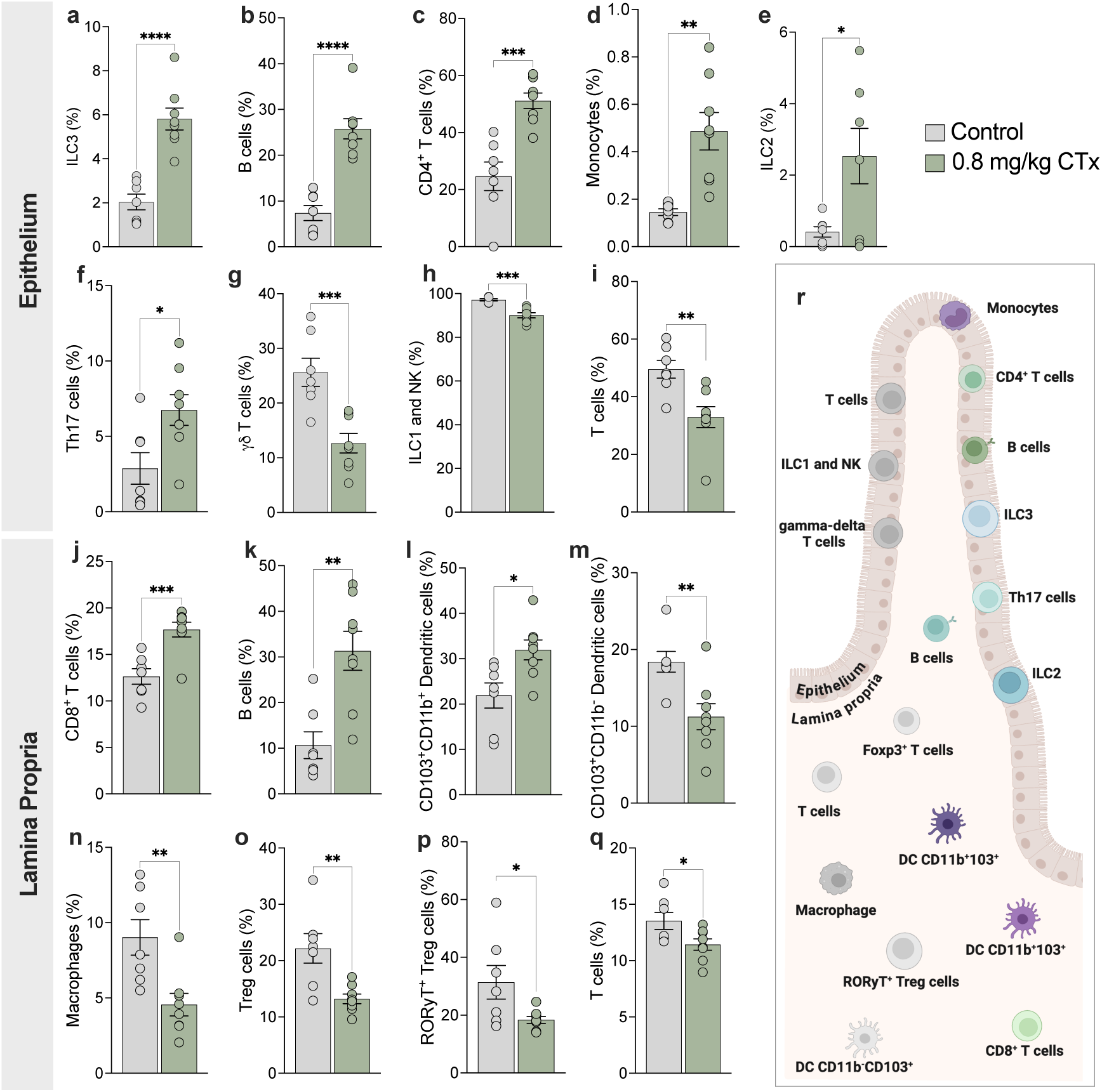
Acute oral CTx exposure remodels immune cell composition in the small intestine. Small intestines from C57BL/6 male mice were collected 24 h after a single oral exposure to cholera toxin (CTx), enzymatically dissociated, and analyzed by full-spectrum flow cytometry (n= 7-8 mice per group). In the intestinal epithelium, CTx altered the frequency of **(a)** ILC3s (CD90.1^+^ TCR-β^−^ TCR-γδ^−^ RorγT^+^), **(b)** B cells (TCR-β^−^ CD19^+^), **(c)** CD4⁺ T cells (CD90.1^+^ TCR-β^+^ CD4^+^), **(d)** monocytes (CD11b^−^ F4/80^−^ Ly6C^+^), **(e)** ILC2s (CD90.1^+^ TCR-β^−^ TCR-γδ^−^ GATA3^+^), **(f)** Th17 cells (CD90.1^+^ TCR-β^+^ CD4^+^ CD44^+^ RorγT^+^), **(g)** γδ T cells (CD90.1^+^ TCR-γδ^+^), **(h)** ILC1/NK cells (CD90.1^+^ TCR-β^−^ TCR-γδ^−^ GATA3^−^ RorγT^−^), **(i)** T cells (CD90.1^+^ TCR-β^+^). In the lamina propria, CTx altered the frequency of **(j)** CD8⁺ T cells (CD90.1^+^ TCR-β^+^ CD8^+^), **(k)** B cells (TCR-β^−^ CD19^+^), **(l)** CD103⁺CD11b⁺ dendritic cells (CD11c^+^MHC-II^+^ CD103^+^ CD11b^+^), **(m)** CD103⁺ dendritic cells (CD11c^+^MHC-II^+^CD11b^−^ CD103^+^), **(n)** macrophages (CD11b^+^ CD64^+^ MHC-II^+^ Ly6C^−^ F4/80^+^), **(o)** regulatory T cells (Treg) cells (CD90.1^+^ TCR-β^+^ CD4^+^ Foxp3^+^), **(p)** RORγt⁺ Treg cells (CD90.1^+^ TCR-β^+^ CD4^+^ Foxp3^+^ RORγT^+^), and **(q)** T cells (CD90.1^+^ TCR-β^+^). **(r)** Summary schematic of flow cytometry findings. All populations were gated on CD45^+^ live cells. Data are presented as mean ± s.e.m. Statistical significance was assessed using unpaired two-tailed *Student’s t*-test. * p ≤ 0.05, ** p≤ 0.01, *** p≤ 0.001, **** p≤ 0.0001. Data are pooled from two independent experiments.

To determine whether this response was sex-dependent, we next examined the same panel in C57BL/6 mice, at 24 h following CTx exposure. The epithelial compartment showed a partially overlapping pattern, with increased frequencies of CD4^+^ T cells and B cells, together with a decreased frequency of total T cells. In the lamina propria, a reduction in the frequency of RORγT⁺ regulatory T cells was the only overlapping finding (Suppl. Figure 5).

Differences in genetic background are known to result in distinct immune responses between C57BL/6 and BALB/c mice in several immune challenge models^30,31^. To assess whether this factor also influenced the response to CTx, we examined intestinal immune cell composition in BALB/c mice of both sexes after CTx exposure. Overall, BALB/c mice exhibited fewer immune changes than C57BL/6 mice, at least for the populations that were analyzed. Among the alterations observed in female BALB/c mice, only the increased frequency of ILC3s and reduced frequency of total T cells in the epithelial compartment, together with the decreased frequency of RORγT⁺ regulatory T cells in the lamina propria, were similar to those observed in male C57BL/6 mice (Suppl. Figure 6). Male BALB/c mice exhibited the fewest changes. The frequencies of ILC3s and ILC1/NK, which were increased and reduced in the epithelial layer, respectively, overlapped with the changes observed in C57BL/6 mice (Suppl. Figure 6).

Together, these analyses identify changes in intestinal immune-cell frequencies 24 h after a single oral CTx exposure. Increased epithelial ILC3 frequencies were detected in C57BL/6 males and in both BALB/c sexes, while reduced epithelial ILC1/NK frequencies were detected in males of both strains. Reduced lamina propria RORγT⁺ Treg frequencies were detected in C57BL/6 males and females and in BALB/c females. These recurring findings occurred alongside other changes specific to the cohorts examined.

### Acute cholera toxin exposure is associated with changes in fecal community composition

We next examined paired fecal samples collected at baseline and 24 h from male C57BL/6 mice receiving CTx (n = 6) or vehicle (n = 5). Within-mouse Bray–Curtis dissimilarity between baseline and 24 h was greater in the CTx group (median, 0.697 versus 0.435; two-sided exact randomization test of the difference in group means, nominal p = 0.045; Suppl. Figure 7a-b). No between-group difference in Jaccard displacement was detected (p = 0.855). Changes in observed ASV richness and Pielou evenness also did not differ significantly between groups (p = 0.361 and p = 0.859, respectively). Thus, the detected community-level signal involved abundance-weighted compositional change; these results do not establish equivalent richness or evenness between treatments. Exploratory inspection identified increases in relative abundance of an Akkermansia-like ASV, Tannerellaceae, and Enterococcaceae in treated mice (Suppl. Figure 7c-e). The latter two patterns were primarily attributable to ASVs assigned to *Parabacteroides goldsteinii* and *Enterococcus faecalis*, respectively. Increased Enterobacteriaceae relative abundance was observed in only a subset of treated mice. These taxon-level patterns are descriptive because taxa were selected after inspection of the data and comparisons were not adjusted for multiple testing.

## DISCUSSION

Although CTx is well recognized for inducing intestinal fluid secretion and promoting mucosal immunity, how its early effects on intestinal tissues relate to whole-body physiology remains poorly understood. The intestine must respond to toxin-induced perturbations while maintaining barrier function, creating potentially competing demands for defense, tissue maintenance, and energy use. Here, we show that a single oral exposure to CTx induces systemic hypometabolism together with intestinal transcriptional and cellular remodeling. Specifically, CTx suppresses whole-body energy expenditure, alters hepatic metabolic gene expression, and induces intestinal signatures associated with defense and maintenance, accompanied by changes in epithelial cells and immune cell composition. In a paired fecal analysis, CTx exposure was also associated with a larger abundance-weighted change in community composition over 24 h.

The physiological response to CTx differed substantially between mouse strains. C57BL/6 mice developed pronounced hypothermia and diarrhea, whereas BALB/c mice exhibited modest temperature changes and intestinal fluid accumulation without diarrhea during the initial observation period. Differences in toxin recognition or processing may contribute to these outcomes. Higher jejunal expression of *B3galt4* in C57BL/6 mice is consistent with greater capacity for GM1 synthesis but does not establish differences in receptor abundance or toxin uptake. CTx can also bind fucosylated glycoproteins and glycolipids independently of GM1^32,33^, and strain differences in gastrointestinal transit or permeability may influence effective toxin exposure^34,35^. Thus, the basis of the distinct physiological responses remains unresolved.

Acute oral CTx exposure induced marked hypothermia in C57BL/6 mice, a response that, to our knowledge, has not previously been reported. Hypothermia also occurs following systemic exposure to LPS or *E. coli* and oral exposure to deoxynivalenol^36–39^. In some inflammatory settings, this response represents a regulated reduction in energy expenditure that limits tissue damage and improves survival^40–42^. However, CTx suppressed metabolism in both C57BL/6 males and BALB/c females despite their distinct thermal responses, indicating that metabolic suppression can occur without pronounced hypothermia. BALB/c mice develop hypothermia following systemic LPS administration^43^, suggesting that their limited response to CTx does not reflect an intrinsic inability to engage thermoregulatory mechanisms. Whether CTx-induced hypothermia reflects regulated energy conservation, fluid and hemodynamic disturbances, or other consequences of intoxication requires further investigation.

The hepatic response to CTx was selective and varied with sex and strain. In C57BL/6 males, CTx increased expression of stress and lipid metabolism genes without coordinated induction of the canonical acute-phase-associated transcripts examined. A shared feature across strains and sexes was suppression of *Cyp7a1*, which encodes the rate-limiting enzyme in classical bile acid synthesis^44^. *Cyp7a1* suppression has also been observed during sustained inflammation and following exposure to other toxicants^26,45–47^, suggesting that altered bile acid metabolism may be a recurrent response to physiological stress. Enteric bacterial infection has been shown to trigger hepatic transcriptional programs that remodel the bile acid metabolome^48^, supporting the idea that gut-derived signals reprogram bile acid metabolism during acute enteric challenge. In C57BL/6 males, increased *Ppara* and *Cpt1a* expression was consistent with altered regulation of fatty acid oxidation and may represent a compensatory response to systemic metabolic suppression. Measurements of bile acid pools and metabolic flux will be needed to establish the functional consequences of these transcriptional changes.

The intestinal transcriptional response suggests concurrent engagement of defense and maintenance programs. Under unfavorable conditions, organisms can reduce investment in growth and prioritize processes that support survival^28^. Consistent with this framework, CTx reduced expression of cell-cycle and DNA-replication genes while inducing genes associated with autophagy, nutrient stress, and lipid utilization. Autophagy can support cellular recycling and stress tolerance, and protective roles have been described in other toxin-exposure models^49–52^. At the same time, CTx increased expression of glycolytic genes and inflammatory mediators, consistent with concurrent inflammatory and glycolytic transcriptional responses. These signatures may reflect distinct or overlapping responses among intestinal cell populations. Their cellular distribution and any contributions to tissue protection remain unresolved.

CTx increased the number of villus DCLK1-positive cells detected by immunostaining and reduced the number of MUC2-positive epithelial cells. Tuft cells detect luminal signals and can release mediators that regulate mucosal immunity, including IL-25, prostaglandins, and acetylcholine^53^. Their established roles in responses to helminths and protozoa raise the possibility that they also participate in toxin sensing or defense^54^. The reduction in MUC2-positive cells requires a different interpretation. Because CTx exposure and *V. cholerae* infection induce mucus secretion^5,55,56^, the reduced intracellular mucin could lower MUC2 immunoreactivity without indicating loss of goblet cells. A recent study in *V. cholerae*-infected infant mice found that IL-22Fc promoted secretory-lineage and MUC2-producing cells^56^, providing evidence for cytokine regulation of this compartment in an infection model. These findings therefore suggest changes in epithelial composition or functional state, whose contributions to the acute response warrant further study.

A single CTx exposure also reorganized immune cell frequencies in the epithelial and lamina propria compartments. Increased epithelial ILC3 frequencies and decreased ILC1/NK frequencies in males of both strains suggest a shift toward RORγt-associated innate populations. Because ILC3 are a major source of IL-22, their increased relative frequency could represent a response to the epithelial changes described above. HIF-1α can promote intestinal ILC3 responses and regulate ILC plasticity^57,58^, providing a possible connection to the intestinal glycolytic signature. However, increased *Hif1a* transcription alone does not establish HIF-1α activation or tissue hypoxia. Intestinal *Il1b* and *Il33* induction could contribute to the observed changes in ILC3 and ILC2 populations, respectively. Notably, IL-33 signaling through its receptor ST2 has previously been shown to be required for robust oral allergic sensitization to peanut in a model using CTx as a mucosal adjuvant, linking this pathway to the development of CTx-facilitated type 2 immunity^16^. The recurrent reduction in RORγt-positive regulatory T-cell frequencies may additionally reflect altered luminal signals, including microbial or bile-acid-derived metabolites that support these cells^59^. These potential connections between tissue metabolism and immune composition remain mechanistic hypotheses.

The paired fecal analysis extends these observations to the intestinal microbial environment. CTx-treated mice showed greater within-mouse Bray–Curtis displacement over 24 h, whereas no corresponding treatment-associated difference was detected using the presence/absence-based Jaccard metric or the reported richness and evenness measures. This pattern is consistent with an abundance-weighted compositional response, but does not establish a common direction of change among treated animals. Changes in secretion, intestinal transit, food intake, or the luminal environment could contribute to the fecal pattern. Because the analysis measured relative abundance in feces, it does not establish changes in absolute bacterial load or in small-intestinal mucosa-associated communities. The taxon-level findings are exploratory. Although altered microbial signals could relate to the immune-frequency changes observed here, the present datasets do not establish that relationship or its direction.

Together, our findings show that acute CTx exposure suppresses whole-body metabolism and is accompanied by intestinal transcriptional, epithelial, immune-cell-frequency, and fecal-community changes. The coexistence of inflammatory and maintenance-associated transcriptional signatures provides a framework for interpreting the early mucosal response. Whether these responses are coordinated and contribute to protection from toxin-induced injury remains to be established.

## LIMITATIONS

Our findings reveal intestinal immune remodeling with systemic hypometabolism after acute CTx exposure, but several questions remain. Bulk jejunal RNA sequencing does not resolve the cellular sources of transcriptional changes and may obscure responses in rare populations. Because transcriptional and cellular analyses were performed at different time points, they do not establish a sequence from early stress signaling to subsequent tissue remodeling. Flow cytometry measures relative frequencies rather than absolute cell numbers and therefore cannot distinguish recruitment, proliferation, redistribution, or cell loss. It also provides incomplete coverage of immune populations and their spatial organization. Also, reduced MUC2-positive counts do not establish goblet-cell loss. The causal relationships among systemic hypometabolism, intestinal transcriptional responses, and immune remodeling remain unresolved. In particular, the defense and maintenance signatures do not establish that these responses protect against toxin-induced injury. Cell-specific analyses and functional perturbations will be needed to determine their contributions to host defense. The metabolic analyses were performed in selected sex–strain combinations under different exposure conditions, limiting direct comparisons between sexes and strains. Contributions of food intake, activity, and fluid balance to the metabolic and thermal phenotypes were not resolved by the results presented here. Finally, this study examines a single acute exposure; the persistence, reversibility, and consequences of the intestinal changes, as well as responses to repeated exposure, remain to be defined. Microbiota profiling was limited to paired baseline and 24-h fecal samples from male C57BL/6 mice. Relative-abundance measurements do not quantify absolute bacterial load or directly characterize the small-intestinal mucosal microbiota. Nonsignificant diversity tests do not establish equivalence, and taxon-level patterns were selected after inspection of the data and analyzed without adjustment for multiple comparisons. The data do not establish whether microbial changes contribute to, or result from, the host response.

## MATERIALS AND METHODS

### Animals

Male and female C57BL/6 and BALB/c mice (6-8 weeks old; Charles River Laboratories, USA), were used in the experiments. All animals were housed in temperature- and humidity-controlled rooms, in a 12/12h light/dark cycle, with food and water provided *ad libitum*. Experiments were conducted in the animal facility at the Biodesign Institute, Arizona State University (Arizona, USA), and Yale University School of Medicine. All animal care and experimentation were approved by the Institutional Animal Care and Use Committee of Arizona State University and Yale University School of Medicine, and consistent with the National Institutes of Health, USA, guidelines.

### Cholera toxin administration

Cholera toxin (CTx, Sigma-Aldrich, 227036) was initially reconstituted in sterile molecular-grade water. CTx was administered intragastrically (i.g) at 0.8 mg/kg or 1.6 mg/kg in 0.2 M sodium carbonate solution (pH 9), in a final volume of 150 μL. Control animals received the same volume of the vehicle solution alone. Oral gavage was performed using plastic needles (Fine Science Tools, 18961-20) during the dark phase, at zeitgeber time 14-15 (ZT14-15).

### Body temperature

Body temperature was measured using a rectal probe (Digi-Sense Model WD 20250-91). Mice were monitored every 30 minutes for up to 6 h post-gavage. A water-based lubricant (Lubricant Liquid, CVS Health) was applied to the probe to minimize irritation, and the probe was sanitized with 70% ethanol between animals.

### Diarrhea and intestinal fluid accumulation

Diarrhea was determined by visual assessment of stool consistency, wetness, and frequency upon oral exposure to CTx and vehicle solution. Intestinal fluid accumulation in the small intestine was scored as present or absent. Fluid and diarrhea analysis were assessed by an observer blinded to group allocation.

### Reduced glutathione (GSH) quantification

Jejunum was collected 6 h after CTx oral gavage, cut longitudinally, and repeatedly washed in phosphate-Buffered Saline (PBS) to remove feces. GSH concentration was determined as previously described^60^ with minor adaptations. Briefly, 150 μL of jejunum supernatant was deproteinized with 2% metaphosphoric acid (1:1) and centrifuged at 7000 g for 10 min at 4°C. Later, 30 μL of deproteinized supernatant was added to a reaction mixture containing 185 μL of 100 mM sodium phosphate buffer (pH 8.0) with 5 mM EDTA, and 15 μL of o-phthaldialdehyde (1 mg/mL in methanol). The mixture was incubated in the dark at room temperature for 15 minutes. The fluorescent product formed by the reaction between GSH and o-phthaldialdehyde was measured using a SpectraMax M5 plate reader (Molecular Devices, Sunnyvale, CA, USA), at excitation and emission wavelengths of 350 and 420 nm, respectively. A calibration curve was generated using a standard GSH solution (0.001–1 mM; Supelco, PHR1359), and results were expressed as nmol GSH/mg protein.

### Protein quantification

Protein content was measured as described by Lowry et al. (1951)^61^. In summary, 10 μL of sample supernatant was incubated in the dark for 10 minutes on a shaker with a reaction mixture containing copper (II) sulfate (Sigma-Aldrich, C1397), potassium sodium tartrate (Sigma-Aldrich, 25508), and 0.1 N NaOH with 2% Na_2_CO_3_ solution (mixed at a 1:1:50 ratio). After incubation, 100 μL of 1N Folin & Ciocalteu’s phenol reagent (Sigma-Aldrich, F9552) was added, and the mixture was incubated for an additional 20 min on a shaker in the dark. Absorbance was measured at 650 nm using a SpectraMax M5 plate reader (Molecular Devices, Sunnyvale, CA, USA). A calibration curve was generated using a standard bovine serum albumin (10–80 μg; Sigma-Aldrich, A7030). Protein concentrations were expressed as mg protein/mL.

### Metabolic assessment

The Promethion High-Definition Metabolic and Behavioral Phenotyping System (Sable Systems International) was used to assess the metabolic profiles (VO_2_, VCO_2_, energy expenditure (EE), and Respiratory Exchange Ratio (RER)) of mice. The mice were placed on a standard 12 h light/12 h dark cycle with a 48-h acclimation period followed by 75 h of sampling (5 h before treatment and 70 h after treatment). Data acquisition and instrument control were executed with Promethion Live Software, and the obtained raw data were processed using ExpeData Data Analysis software. These experiments were performed at Yale School of Medicine. For each metabolic parameter, the area under the curve (AUC) was calculated per mouse over the interval from 65 to 1,000 min (approximately 1 to 16.7 h) after CTx or vehicle administration for C57BL/6 mice, and from 185 to 1,000 min (approximately 3 to 16.7 h) for BALB/c mice. Group means were then compared, and the percentage change in CTx-treated mice was expressed relative to the control mean (set as 100%).

### Tissue digestion and cell isolation

Enzymatic digestion of the small intestine was performed as described previously^14^ with minor modifications. Following harvest, the small intestine was collected by separating it distal to the pyloric valve and proximal to the cecum. Fat tissue and Peyer’s Patches were removed and discarded. The intestine was cut open longitudinally and repeatedly washed in 1x Phosphate-Buffered Saline (PBS) to remove fecal matter. After cleaning, each small intestine was cut into 2 cm pieces and placed into an Erlenmeyer flask containing room-temperature DMEM supplemented with 3% FBS, 5 mM EDTA, and 0.94 mM dithiothreitol. Samples were incubated with stirring (600 rpm) at 37°C, 5% CO_2_ for 20 min. The pieces were then washed three times by vigorous shaking (30 s each) in FBS-free DMEM containing 2 mM EDTA. After each wash, the medium was passed through a strainer and collected in a beaker on ice to obtain the epithelial fractions.

#### Lamina Propria fraction

The intestinal tissue remaining in the strainer was finely minced and incubated with stirring (∼ 600 rpm) at 37 °C and 5% CO_2_ in 10 mL FBS-free DMEM containing 0.1 mg/mL Liberase TL (Sigma-Aldrich, 5401020001) and 0.05% Dnase I (Sigma-Aldrich, D5025) for 30 min. The digestion was stopped by adding 10 mL cold DMEM with 3% FBS, followed by washing the beaker with an additional 20 mL cold DMEM with 3% FBS. The sample was then passed through a 70 μm cell strainer using the plunger of a 3 mL syringe and centrifuged at 268 x g for 5 min at 4°C. The pellet was resuspended, filtered again through a 40 μm cell strainer, and centrifuged under the same conditions. The resulting pellet contained lamina propria immune cells.

#### Epithelial fraction

The cell suspension collected on ice during the washes was filtered through a 70 μm cell strainer and centrifuged at 268 x g, for 8 min at 4°C. The pellet was resuspended, filtered again through a 40 μm cell strainer, and centrifuged under the same conditions. The final pellet contained the epithelial immune cell fraction.

### Flow cytometry

Small intestines were collected 24 h after CTx oral gavage. The epithelial and lamina propria cell fractions were obtained as described above. Cell suspensions (0.75-1 × 10^6^ cells/ sample) were transferred to U-bottom 96-well plates, washed twice with assay buffer (2% BSA in PBS) by centrifugation (5 min, 4°C, 268 x g). Cell populations were stained with Fc block (anti-CD16/32) and fluorochrome-conjugated antibody cocktail (**Table 1**) in assay buffer, targeting cell surface markers on ice for 30 min. After surface staining, cells were fixed in 1% PFA (in PBS) at room temperature for 20 min.

**Table 1.**
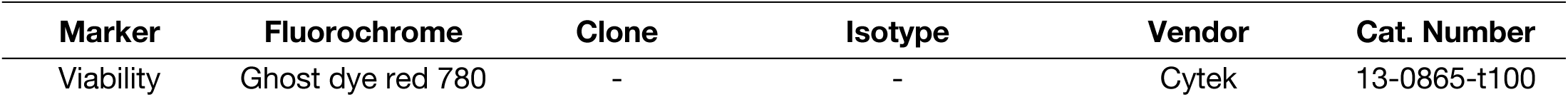
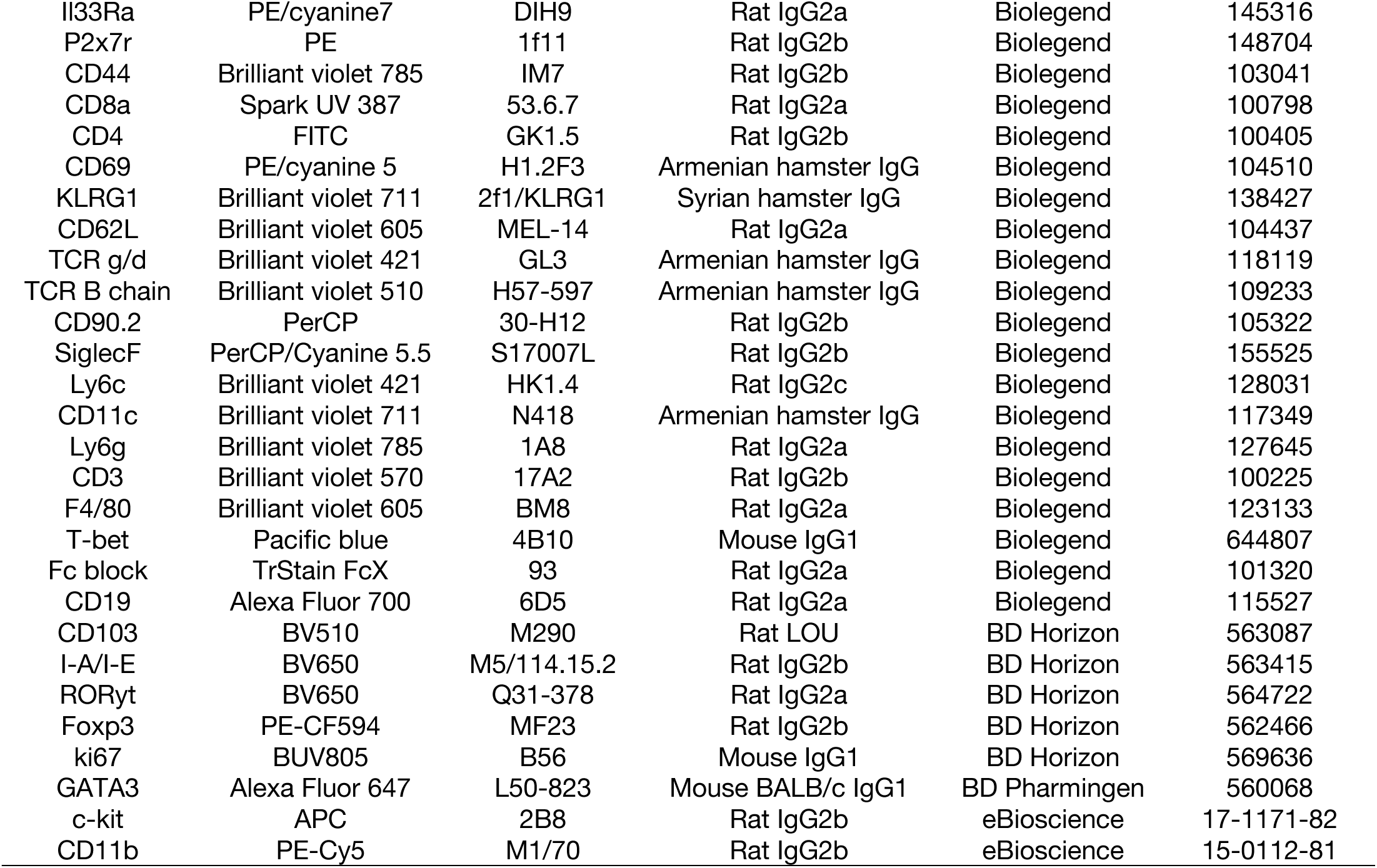
Antibodies list used for Full-spectrum flow cytometry analysis.

For intracellular staining, cells were first stained for surface epitopes, fixed, and subsequently permeabilized (eBioscience, 00-5123-43) before overnight staining according to the manufacturer’s protocol. The gating strategy used was as follows: all analyzed cells were gated on live CD45^+^ events. <u>Neutrophils</u>: CD11b^+^ Ly6G^+^, <u>monocytes</u>: CD11b^+^ CD64^−^ MHC-II^−^ F4/80^−^ Ly6C^+^, e<u>osinophils</u>: CD11b^+^ Siglec-F^+^ MHC-II^−^ CD11c^−^ Ly6C^−^, <u>macrophages</u>: CD11b^+^ CD64^+^ MHC-II^+/-^ F4/80^+^ Ly6C^−^, <u>dendritic cells</u>: CD11c^+^ MHC-II^+^ CD19^−^ TCR-β^−^ SiglecF^−^ Ly6G^−^ Ly6C^−^ CD64^−^ CD103^+^ or CD11b^+^, <u>B-cells</u>: TCR-β^−^ CD19^+^, <u>T-cells</u>: CD90.2^+^ TCR-β^+^, CD8^+^ T cells: CD90.2^+^ TCR-β^+^ CD8^+^, CD4^+^ T cells: CD90.2^+^ TCR-β^+^ CD4^+^, <u>ILC1/NK cells</u>: CD90.2^+^ TCR-β^−^ TCR-γδ^−^ GATA-3^−^ RORγT^−^ Tbet^+^, <u>ILC2</u>: CD90.2^+^ TCR-β^−^ TCR-γδ^−^ GATA-3^+^ RORγT^−^, <u>ILC3</u>: CD90.2^+^ TCR-β^−^ TCR-γδ^−^ GATA-3^−^ RORγT^+^, <u>Th1 cells</u>: CD90.2^+^ TCR-β^+^ CD4^+^ CD8^−^ Tbet^+^ GATA-3^−^ RORγT^−^ FoxP3^−^, <u>Th2</u> <u>cells</u>: CD90.2^+^ TCR-β^+^ CD4^+^ CD8^−^ Tbet ^−^ GATA-3^+^ RORγT^−^ FoxP3^−^, <u>Th17 cells</u>: CD90.2^+^ TCR-β^+^ CD4^+^ CD8^−^ Tbet ^−^ GATA-3^−^ RORγT^+^ FoxP3^−^, <u>Regulatory T cells (Treg)</u>: CD90.2^+^ TCR-β^+^ CD4^+^ CD44^+^ GATA-3^+/-^ RORγT^+/-^ FoxP3^+^, and <u>γδ T-cells</u>: CD90.2^+^ TCR-γδ^+^ TCR-β^−^.

Full-spectrum flow cytometry was performed using a Cytek Aurora (Cytek Biosciences), at the Department of Immunology, Mayo Clinic, Phoenix, AZ. Data were analyzed with FlowJo Software (Version 10.8.0 BD Biosciences).

### RNA extraction and quantification

Liver and jejunum samples were collected 6 h after CTx oral gavage. For RNA extraction, tissues were harvested in RNA STAT-60 RNA isolation reagent (Amsbio, CS-502-500) and homogenized by ceramic bead homogenization (Omni, INC). RNA was extracted using the Rneasy Mini Kit (Qiagen, 74106) for liver samples and Direct-zol RNA Mini Prep (ZYMO Research, R2072) for small intestine samples. cDNA synthesis was performed using SMART^®^ MMLV reverse transcriptase (Takara, 639524) with oligo(dT) primers. Quantitative real-time PCR was conducted on a ViiA 7 Real-Time PCR System (Life Technologies^TM^ – Applied Biosystems®) using Power SYBR™ Green PCR Master Mix (Life Technologies, 4367660). Transcript levels were normalized to *Rpl13a* expression (housekeeping).

**Table 2.**
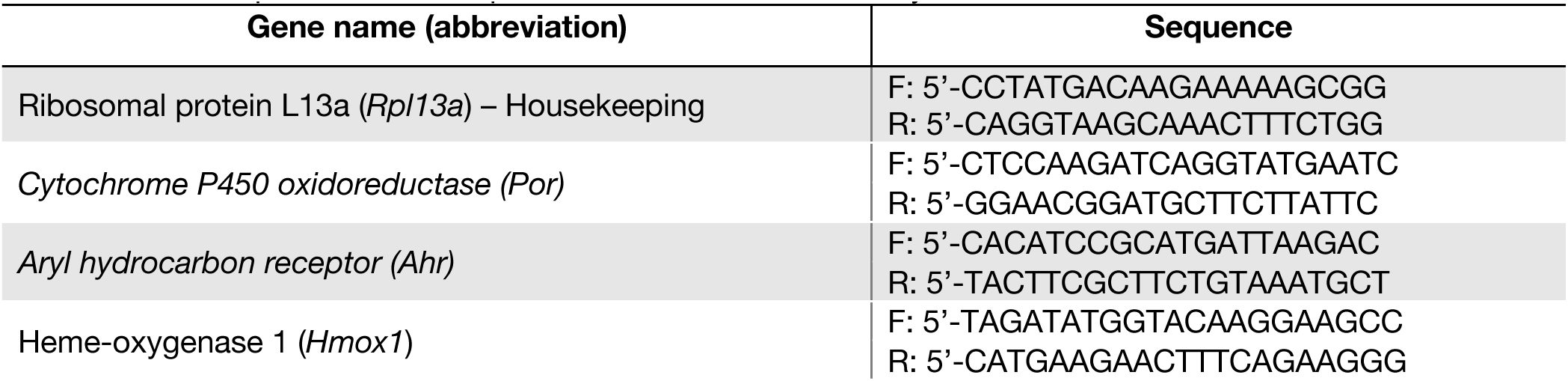
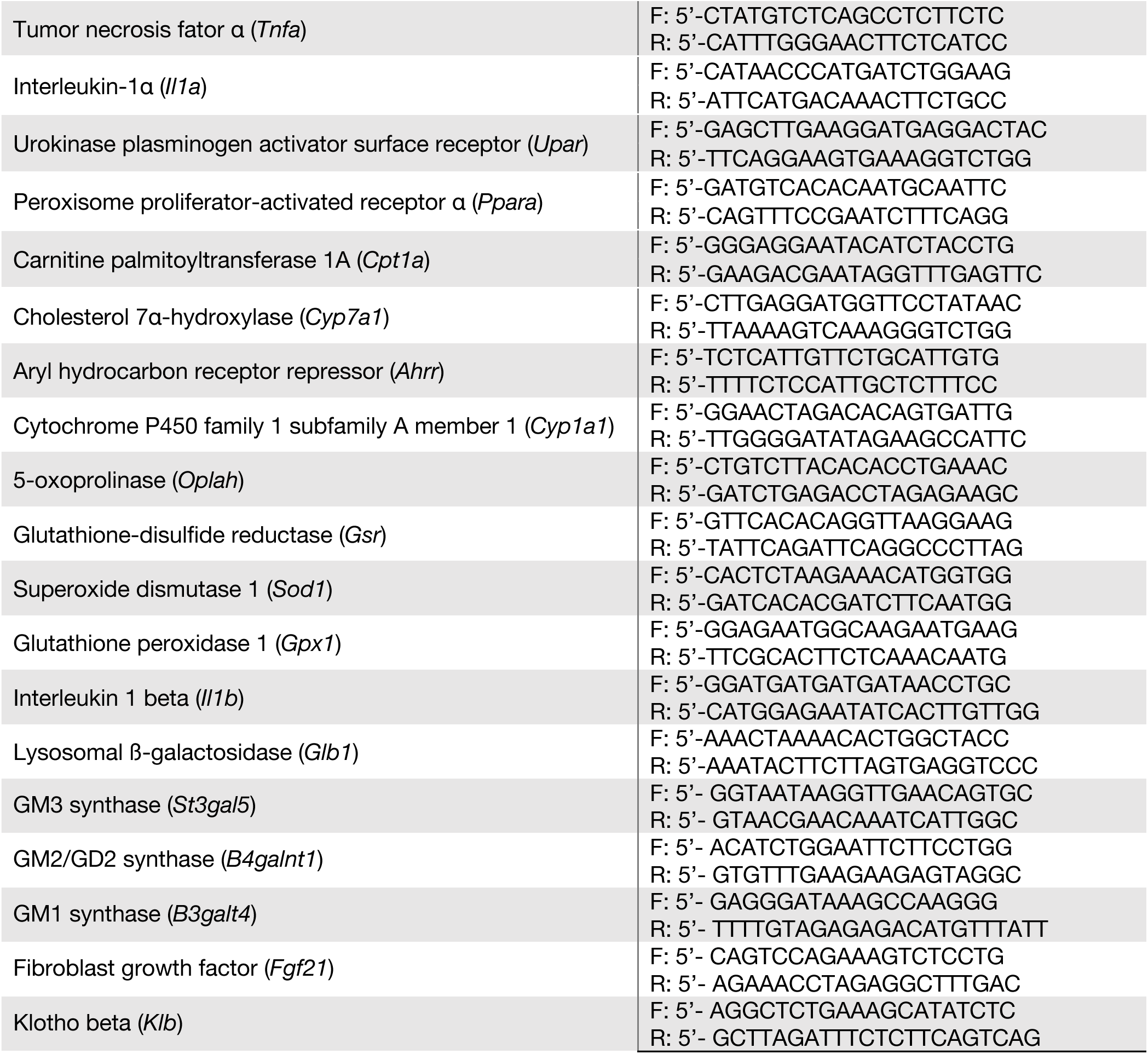
Gene sequence used for quantitative real-time PCR analysis.

### RNA sequencing and data processing

Bulk RNA was extracted from jejunum samples (6 replicates per group) as described above. RNA quality control, library preparation, and sequencing were conducted at Azenta Life Sciences (South Plainfield, NJ, USA) as follows:

Total RNA samples were quantified using Qubit 4.0 Fluorometer (Life Technologies, Carlsbad, CA, USA), and RNA integrity was checked using Agilent TapeStation 4200 (Agilent Technologies, Palo Alto, CA, USA).

RNA sequencing libraries were prepared using the NEBNext Ultra II RNA Library Prep Kit for Illumina using the manufacturer’s instructions (NEB, Ipswich, MA, USA). Briefly, mRNAs were initially enriched with Oligo(dT) beads. Enriched mRNAs were fragmented for 15 minutes at 94 °C. First-strand and second-strand cDNA were subsequently synthesized. cDNA fragments were end-repaired and adenylated at 3’ends, and universal adapters were ligated to cDNA fragments, followed by index addition and library enrichment by PCR with limited cycles. The sequencing library was validated on the Agilent TapeStation (Agilent Technologies, Palo Alto, CA, USA) and quantified using the Qubit 4.0 Fluorometer (Invitrogen, Carlsbad, CA) and quantitative PCR (KAPA Biosystems, Wilmington, MA, USA).

The sequencing libraries were multiplexed and clustered onto a flowcell on the Illumina NovaSeq instrument according to the manufacturer’s instructions. The samples were sequenced using a 2×150bp Paired End (PE) configuration. Image analysis and base calling were conducted by the NovaSeq Control Software (NCS). Raw sequence data (.bcl files) generated from Illumina NovaSeq were converted into fastq files and de-multiplexed using Illumina bcl2fastq 2.20 software. One mismatch was allowed for index sequence identification.

Raw sequencing data were evaluated for quality using FastQC (v0.12.1). Adapter sequences were removed using Fastp (v0.23.4) with default parameters. High-quality reads were aligned to the mouse reference genome (GRCm39/mm39.114) using STAR (v2.7.11a) with default settings optimized for paired-end reads. Gene-level read counts were quantified using Stringtie (v2.2.3), based on the GENCODE M34 annotation. Downstream analyses, including differential gene expression, were performed using DESeq2 (v1.42.0) in R.

The RNA-sequencing data generated in this study has been deposited in the NCBI Gene Expression Omnibus (GEO).

### Histology and Immunofluorescence Assay

Specialized intestinal epithelial cells were quantified by immunofluorescence staining. Briefly, the proximal portions of the jejunum, approximately 6cm, were collected 24 h post-exposure. Sections were flushed with 1x PBS for removal of fecal matter, and were then opened longitudinally and subsequently rolled into Swiss rolls and fixed overnight 4% paraformaldehyde at 4°C. The intestines were then dehydrated, embedded in paraffin, and cut into 5 μm sections.

Antigen retrieval was performed using Borg Decloaker RTU solution (Biocare Medical, BD1000G1) and a pressurized Decloaking Chamber (BioSB, TintoRetriever Pressure Cooker). The following primary antibodies were used for immunofluorescence assay: Rabbit polyclonal anti-Muc2 (1:500, Novus, NBP1-31231), Rabbit polyclonal anti-Lyz1 (1:250, FisherSci, RB372A1), Rabbit monoclonal Anti-DCLK1 (1:250, Abcam ab109029), or Rabbit Anti-Chromogranin A (1:250, Abcam, 15160). Nuclear staining was performed using DAPI (1:1000, Biolegend 422801). The Alexa Fluor secondary antibody Anti-Rabbit 568 (1:500, Invitrogen, A10042) was used for visualization of all the primary antibodies listed above. All Primary and Secondary antibody dilutions were performed in SignalStain Antibody Diluent (CST, 8112L). Tissue was mounted using Invitrogen Prolong Gold mounting medium. Images were acquired using a 20x objective (Olympus VS200 ASW) using an Olympus SlideView VS200 upright microscope.

### Fecal sampling, DNA extraction, and full-length 16S rRNA gene sequencing

Freshly voided fecal pellets were collected from male C57BL/6 mice immediately before and 24 h after intragastric gavage with vehicle (n= 5) or 0.8 mg/kg CTx (n= 6), giving 22 samples comprising 11 baseline–24-h pairs. Each mouse was placed in a separate sterilized container for collection; pellets were transferred to microcentrifuge tubes, placed on dry ice, and stored at −70°C. DNA extraction and sequencing were performed by SeqCenter (Pittsburgh, PA, USA). DNA was extracted using the ZymoBIOMICS DNA Miniprep Kit according to the manufacturer’s instructions, including bead-beating lysis (MP FastPrep-24, five cycles) and Zymo-Spin III-HRC inhibitor removal, and quantified by Qubit. Blank, negative, and positive controls were included throughout extraction and sequencing. Full-length (V1-V9) 16S rRNA gene libraries were prepared with the Quick-16S Full-Length Library Prep Kit using 27F and 1492R primers, then prepared for sequencing with the PacBio HiFi Prep Kit 96 and sequenced on a PacBio Revio instrument (12-h run). Circular consensus sequences were generated on instrument and demultiplexed with Lima v2.12.0 (minimum score 80, minimum end score 50, minimum reference span 0.75, two scoring regions); BAM files were converted to FASTQ using SAMtools v1.17.

### Bioinformatic processing and taxonomic assignment

Processing followed the PacBio HiFi-16S-Workflow in QIIME 2 2024.10. Primers were removed with cutadapt and reads denoised with DADA2 to generate an amplicon sequence variant (ASV) table. ASVs were classified with VSEARCH against SILVA 138, GTDB release 207, and Greengenes2. One abundant ASV that was unresolved in the primary taxonomy assignment was classified as *Akkermansia* by Greengenes2. Because its assignment differed across databases, we refer to it as an *Akkermansia*-like ASV. The final ASV table contained 1,350 ASVs and 463,543 reads across 22 samples. All samples exceeded 5,000 reads (range 8,764-22,255) and were retained. Taxon-level analyses were restricted to annotated ASVs.

### Diversity and statistical analyses

Analyses were performed in R 4.5.3 using tidyverse (V1.3.2), vegan (v2.7-6), and ape (v5.8-1). Relative abundance was calculated per sample as each ASV count divided by total sample reads. Alpha diversity was summarized as observed ASV richness, Shannon diversity, inverse Simpson diversity, and Pielou evenness, with analytical rarefaction to the minimum depth (8,764 reads) as a sensitivity analysis. Beta diversity was assessed by Bray-Curtis dissimilarity (relative abundances) and Jaccard distance (presence/absence). The mouse was the experimental unit. Baseline and 24-h samples were paired by mouse, and each mouse’s change in alpha diversity, beta-diversity displacement, or taxon relative abundance was calculated before groups were compared; taxon changes are reported as 24-h minus baseline relative abundance in percentage points. Group differences were assessed by two-sided exact randomization tests of the difference in group means, evaluating all 462 possible label assignments while retaining the observed group sizes. Exploratory taxon-level analyses focused on the *Akkermansia*-like ASV and on Tannerellaceae, Enterococcaceae, Enterobacteriaceae, and Lactobacillaceae, selected on the magnitude and recurrence of within-mouse changes; the Tannerellaceae and Enterococcaceae signals were driven primarily by ASVs assigned to *Parabacteroides goldsteinii* and *Enterococcus faecalis*. Significance was defined as p < 0.05. Taxon-level analyses were exploratory, and their unadjusted p values are descriptive. No microbiota observations were excluded as outliers.

### Statistics analysis

Data normality was assessed using the Shapiro-Wilk test (α=0.05). Depending on distribution, comparisons between two groups were performed using an unpaired t-test (parametric) or the Mann-Whitney test (nonparametric). For multiple group comparisons, an ordinary one-way ANOVA followed by Tukey’s multiple comparisons test (parametric) or a Kruskal-Wallis test (nonparametric) was used. Results were considered statistically significant at p < 0.05. Outliers were identified using the ROUT method in GraphPad Prism (Q = 1%). Data are presented as mean±S.E.M.

## ACKNOWLEDGMENTS

We would like to thank all current and former members of the Florsheim Lab for helpful discussions. We thank the Department of Animal Care and Technologies team for technical assistance and the staff of the Biodesign Institute for their continued support. We are grateful to Jaime Cullen and Ruslan Medzhitov of Yale School of Medicine, for providing the animals and reagents used in the metabolic assessment experiments. We also thank Kristina Buss of the Desert Genome Center Bioinformatics Core at Arizona State University for assistance with coding for the RNA-sequencing analysis. Schematics were created with BioRender.com.

## AUTHOR CONTRIBUTIONS

ACR: Conceptualization, Data curation, Investigation, Methodology, Software, Validation, Visualization, Writing – original draft, Writing – review & editing. MG, CLS, GL: Formal analysis, Investigation. CW: Investigation and Software. THM: Investigation. BGCL: Formal analysis. GPRC, LEVG: Formal analysis, Investigation. HBdS, MOD, MDM: Resources, Writing – review & editing. EBF: Conceptualization, Funding acquisition, Methodology, Project administration, Supervision, Validation, Writing – original draft, Writing – review & editing.

## DECLARATION OF GENERATIVE AI AND AI-ASSISTED TECHNOLOGIES

During preparation of this manuscript, the authors used ChatGPT (OpenAI) to improve conciseness, grammar, and readability, and Consensus for literature search. All suggested revisions were reviewed and edited by the authors, who take full responsibility for the final content of the manuscript.

## REFERENCES

1. Xie, Z. et al. The gut-to-brain axis for toxin-induced defensive responses. Cell 185, 4298–4316.e21 (2022).

2. Song, W. et al. T-2 toxin metabolism and its hepatotoxicity: New insights on the molecular mechanism and detoxification. Environ. Pollut. 330, 121784 (2023).

3. Jobling, M. G., Yang, Z., Kam, W. R., Lencer, W. I. & Holmes, R. K. A Single Native Ganglioside GM1-Binding Site Is Sufficient for Cholera Toxin To Bind to Cells and Complete the Intoxication Pathway. mBio 3, 10.1128/mbio.00401-12 (2012).

4. Merritt, E. A. et al. Crystal structure of cholera toxin B-pentamer bound to receptor GM1 pentasaccharide. Protein Sci. 3, 166–175 (1994).

5. Lencer, W. I., Hirst, T. R. & Holmes, R. K. Membrane traffic and the cellular uptake of cholera toxin. Biochim. Biophys. Acta (BBA) - Mol. Cell Res. 1450, 177–190 (1999).

6. Cong, Y., Oliver, A. O. & Elson, C. O. Effects of cholera toxin on macrophage production of co-stimulatory cytokines. Eur. J. Immunol. 31, 64–71 (2001).

7. Bromander, A. K., Kjerrulf, M., Holmgren, J. & Lycke, N. Cholera Toxin Enhances Alloantigen Presentation by Cultured Intestinal Epithelial Cells. Scand. J. Immunol. 37, 452–458 (1993).

8. Lavelle, E. C. et al. Cholera Toxin Promotes the Induction of Regulatory T Cells Specific for Bystander Antigens by Modulating Dendritic Cell Activation. J. Immunol. 171, 2384–2392 (2003).

9. Blázquez, A. B. & Berin, M. C. Gastrointestinal Dendritic Cells Promote Th2 Skewing via OX40L. J. Immunol. 180, 4441–4450 (2008).

10. Anjuère, F., et al. In Vivo Adjuvant-Induced Mobilization and Maturation of Gut Dendritic Cells after Oral Administration of Cholera Toxin. J. Immunol. 173, 5103–5111 (2004).

11. Marinaro, M. et al. Mucosal adjuvant effect of cholera toxin in mice results from induction of T helper 2 (Th2) cells and IL-4. J. Immunol. 155, 4621–4629 (1995).

12. Braun, M. C., He, J., Wu, C.-Y. & Kelsall, B. L. Cholera Toxin Suppresses Interleukin (IL)-12 Production and IL-12 Receptor β1 and β2 Chain Expression. J. Exp. Med. 189, 541–552 (1999).

13. Lycke, N., Bromander, A. K., Ekman, L., Karlsson, U. & Holmgren, J. Cellular basis of immunomodulation by cholera toxin in vitro with possible association to the adjuvant function in vivo. J. Immunol. 142, 20–27 (1989).

14. Zhang, B. et al. Metabolic fitness of IgA+ plasma cells in the gut requires DOCK8. Mucosal Immunol. 17, 431–449 (2024).

15. Snider, D. P., Marshall, J. S., Perdue, M. H. & Liang, H. Production of IgE antibody and allergic sensitization of intestinal and peripheral tissues after oral immunization with protein Ag and cholera toxin. J Immunol 153, 647–57 (1994).

16. Chu, D. K. et al. IL-33, but not thymic stromal lymphopoietin or IL-25, is central to mite and peanut allergic sensitization. J. Allergy Clin. Immunol. 131, 187–200.e8 (2013).

17. Baselski, V., Briggs, R. & Parker, C. Intestinal fluid accumulation induced by oral challenge with Vibrio cholerae or cholera toxin in infant mice. Infect. Immun. 15, 704–712 (1977).

18. Tang, L.-Q. et al. Calcium/calcimimetic via calcium-sensing receptor ameliorates cholera toxin-induced secretory diarrhea in mice. World J. Gastroenterol. 30, 268–279 (2024).

19. Labarrere, C. A. & Kassab, G. S. Glutathione: A Samsonian life-sustaining small molecule that protects against oxidative stress, ageing and damaging inflammation. Front. Nutr. 9, 1007816 (2022).

20. Lee, M.-L., Chang, C.-P., Toda, C., Nemoto, T. & Enoki, R. Body temperature regulates glucose metabolism and torpid behavior. Nat. Commun. 16, 6278 (2025).

21. Tattersall, G. J. et al. Coping with Thermal Challenges: Physiological Adaptations to Environmental Temperatures. Compr. Physiol. 2, 2151–2202 (2012).

22. Preiser, J.-C., Ichai, C., Orban, J.-C. & Groeneveld, A. B. J. Metabolic response to the stress of critical illness. Br. J. Anaesth. 113, 945–954 (2014).

23. Schuler, C. B., et al. Energy Allocation Resilience and Endocrine Integration. Int. J. Mol. Sci. 27, 1345 (2026).

24. Rui, L. Energy Metabolism in the Liver. Compr. Physiol. 4, 177–197 (2014).

25. Robinson, M. W., Harmon, C. & O’Farrelly, C. Liver immunology and its role in inflammation and homeostasis. Cell Mol Immunol 13, 267–276 (2016).

26. Okin, D. & Medzhitov, R. The Effect of Sustained Inflammation on Hepatic Mevalonate Pathway Results in Hyperglycemia. Cell 165, 343–356 (2016).

27. Huen, S. C. et al. Hepatic FGF21 preserves thermoregulation and cardiovascular function during bacterial inflammation. J. Exp. Med. 218, e20202151 (2021).

28. Wang, A., Luan, H. H. & Medzhitov, R. An evolutionary perspective on immunometabolism. Science 363, (2019).

29. Williamson, E., Westrich, G. M. & Viney, J. L. Modulating Dendritic Cells to Optimize Mucosal Immunization Protocols. J. Immunol. 163, 3668–3675 (1999).

30. Wakeham, J., Wang, J. & Xing, Z. Genetically Determined Disparate Innate and Adaptive Cell-Mediated Immune Responses to Pulmonary Mycobacterium bovis BCG Infection in C57BL/6 and BALB/c Mice. Infect. Immun. 68, 6946–6953 (2000).

31. Fukushima, A. et al. Genetic background determines susceptibility to experimental immune-mediated blepharoconjunctivitis: Comparison of Balb/c and C57BL/6 mice. Exp. Eye Res. 82, 210–218 (2006).

32. Wands, A. M. et al. Fucosylation and protein glycosylation create functional receptors for cholera toxin. eLife 4, e09545 (2015).

33. Cervin, J. et al. GM1 ganglioside-independent intoxication by Cholera toxin. PLoS Pathog. 14, e1006862 (2018).

34. Gama, L. A. et al. Physiological effects of environmental enrichment on BALB/c and C57BL/6 strain mice. Discov. Anim. 2, 7 (2025).

35. Hoyt, L. R. et al. Cysteinyl leukotrienes stimulate gut absorption of food allergens to promote anaphylaxis in mice. Science 389, eadp0240–eadp0240 (2025).

36. Greer, G. G. & Rietschel, E. T. Lipid A-induced tolerance and hyperreactivity to hypothermia in mice. Infect. Immun. 19, 357–368 (1978).

37. Rudaya, A. Y., Steiner, A. A., Robbins, J. R., Dragic, A. S. & Romanovsky, A. A. Thermoregulatory responses to lipopolysaccharide in the mouse: dependence on the dose and ambient temperature. *Am. J. Physiol.-Regul.*, Integr. Comp. Physiol. 289, R1244–R1252 (2005).

38. Liu, E. et al. Naturally occurring hypothermia is more advantageous than fever in severe forms of lipopolysaccharide- and Escherichia coli-induced systemic inflammation. *Am. J. Physiol.-Regul.*, Integr. Comp. Physiol. 302, R1372–R1383 (2012).

39. Girardet, C. et al. Central Inflammation and Sickness-Like Behavior Induced by the Food Contaminant Deoxynivalenol: A PGE2-Independent Mechanism. Toxicol Sci 124, 179–191 (2011).

40. Corrigan, J. J., Fonseca, M. T., Flatow, E. A., Lewis, K. & Steiner, A. A. Hypometabolism and hypothermia in the rat model of endotoxic shock: independence of circulatory hypoxia. J. Physiol. 592, 3901–3916 (2014).

41. Tinajero, A. et al. Spontaneous and pharmacologically induced hypothermia protect mice against endotoxic shock. Br. J. Pharmacol. 182, 2621–2641 (2025).

42. Wanner, S. P. et al. Cold-Induced Thermogenesis and Inflammation-Associated Cold-Seeking Behavior Are Represented by Different Dorsomedial Hypothalamic Sites: A Three-Dimensional Functional Topography Study in Conscious Rats. J. Neurosci. 37, 6956–6971 (2017).

43. Kher, P. et al. Systemic endotoxemia induces integrated sickness physiology in female BALB/c mice. bioRxiv 2026.08.22.746462 (2026).doi:10.64898/2026.08.22.746462

44. Chiang, J. Y. L. & Ferrell, J. M. Up to date on cholesterol 7 alpha-hydroxylase (CYP7A1) in bile acid synthesis. Liver Res. 4, 47–63 (2020).

45. Hari, A. et al. Exposure to PFAS chemicals induces sex-dependent alterations in key rate-limiting steps of lipid metabolism in liver steatosis. Front. Toxicol. 6, 1390196 (2024).

46. Choi, S.-Y. et al. Transcriptomic alterations induced by aflatoxin B1 and ochratoxin A in LMH cell line. Poult. Sci. 99, 5265–5274 (2020).

47. Liu, X. et al. Cyp7a1 and Cyp8b1 Downregulation Characterizes Concanavalin-A-Induced Acute Liver Injury: Insights from Multiomics Analysis. J. Proteome Res. 25, 672–683 (2026).

48. Zhang, T., Hasegawa, Y. & Waldor, M. K. Enteric bacterial infection stimulates remodelling of bile metabolites to promote intestinal homeostasis. Nat. Microbiol. 9, 3376–3390 (2024).

49. Wu, J. et al. Autophagy and Apoptosis Interact to Modulate T-2 Toxin-Induced Toxicity in Liver Cells. Toxins 11, 45 (2019).

50. Wu, Y. et al. Autophagic Degradation Is Involved in Cell Protection against Ricin Toxin. Toxins 15, 304 (2023).

51. Zhao, Y., Lin, H., Ma, J. & Cui, Y. Botulinum toxin type A activates protective autophagy by modulating endoplasmic reticulum stress in hypoxia/reoxygenation-treated endothelial cells. Heliyon 9, e17907 (2023).

52. Tang, Y. et al. Autophagy protects intestinal epithelial Cells against Deoxynivalenol toxicity by alleviating oxidative stress via IKK signaling pathway. Free Radic. Biol. Med. 89, 944–951 (2015).

53. Schneider, C., O’Leary, C. E. & Locksley, R. M. Regulation of immune responses by tuft cells. Nat. Rev. Immunol. 19, 584–593 (2019).

54. Hendel, S. K. et al. Tuft Cells and Their Role in Intestinal Diseases. Front. Immunol. 13, 822867 (2022).

55. Forstner, J. F., Roomi, N. W., Fahim, R. E. & Forstner, G. G. Cholera toxin stimulates secretion of immunoreactive intestinal mucin. Am. J. Physiol.-Gastrointest. Liver Physiol. 240, G10–G16 (1981).

56. Suzuki, M. et al. IL-22 promotes genesis of small intestinal secretory cells that protect against cholera in mice. Nat. Microbiol. 11, 1949–1966 (2026).

57. Fachi, J. L. et al. Hypoxia enhances ILC3 responses through HIF-1α-dependent mechanism. Mucosal Immunol. 14, 828–841 (2021).

58. Krzywinska, E. et al. The transcription factor HIF-1α mediates plasticity of NKp46+ innate lymphoid cells in the gut. J. Exp. Med. 219, e20210909 (2022).

59. Song, X. et al. Microbial bile acid metabolites modulate gut RORγ+ regulatory T cell homeostasis. Nature 577, 410–415 (2020).

60. Browne, R. W. & Armstrong, D. Free Radical and Antioxidant Protocols. Methods Mol. Biol. (Clifton, NJ) 108, 347–352 (1998).

61. Lowry, O. H., Rosebrough, N. J., Farr, A. L. & Randall, R. J. Protein measurement with the Folin phenol reagent. J. Biol. Chem. 193, 265–75 (1951).

