## Supplemental Figure 1 for "Cholera toxin remodels intestinal immunity and suppresses systemic metabolism"

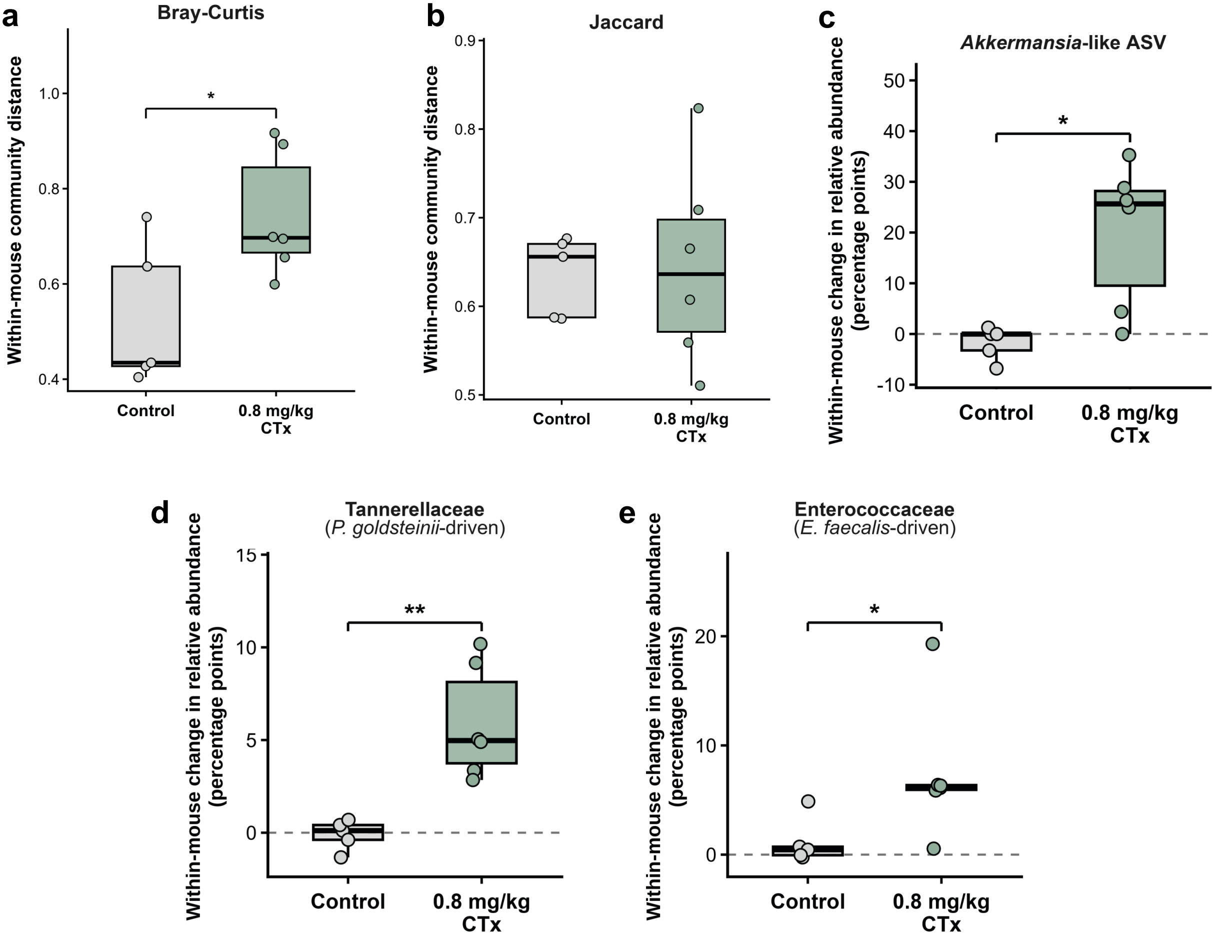


**Supplementary Figure 7 | Acute oral CTx exposure is associated with fecal microbial community restructuring.** Within-mouse Bray-Curtis and Jaccard distances quantify the change between each mouse's baseline and 24-h microbial community. **(a)** Bray-Curtis dissimilarity was calculated from ASV relative abundances, and **(b)** Jaccard distance was calculated from ASV presence or absence. Bray-Curtis displacement was greater following CTx exposure ($p = 0.045$), whereas Jaccard displacement did not differ between groups ($p = 0.855$). Within-mouse changes in relative abundance are shown for an **(c)** *Akkermansia*-like ASV, **(d)** Tannerellaceae (*P. goldsteinii*-driven), and **(e)** Enterococcaceae (*E. faecalis*-driven). Significant treatment-associated changes were observed for the *Akkermansia*-like ASV ($p = 0.022$), Tannerellaceae ($p = 0.004$), and Enterococcaceae ($p = 0.017$) between groups. The *Akkermansia*-like feature corresponds to ASV d5ebbbf0f104adc463156fca557bb6fc, which was unresolved below Bacteria by GTDB; its display label reflects the secondary GreenGenes2 assignment. Taxon panels use separate y-axis ranges. Each point represents one mouse. Boxes show the median and interquartile range, and whiskers extend to 1.5 times the interquartile range. Group comparisons were performed using two-sided exact randomization tests of the mean within-mouse change or distance, enumerating all 462 possible assignments of six CTx labels among 11 mice. Only statistically significant comparisons are marked. * $p \leq 0.05$; ** $p \leq 0.01$. Taxon-level comparisons were exploratory and were not adjusted for multiple comparisons.
