## Supplemental Figure 2 for "Cholera toxin remodels intestinal immunity and suppresses systemic metabolism"

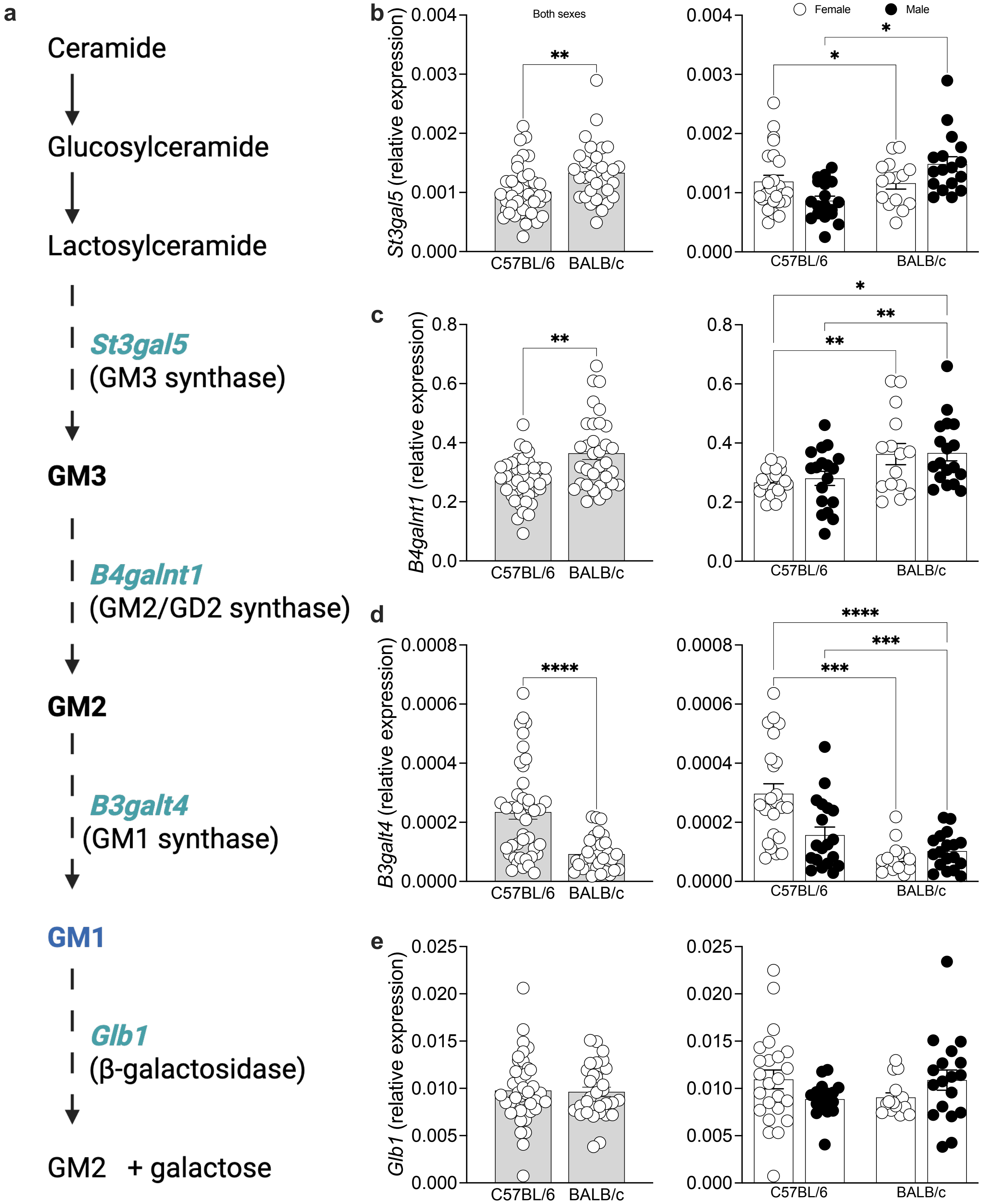


**Supplementary Figure 1 | Intestinal GM1 ganglioside receptor pathway gene expression across mouse strains and sexes.** Jejunum samples from C57BL/6 and BALB/c mice of both sexes were analyzed by quantitative real-time PCR for key genes involved in the GM1 ganglioside synthesis and metabolism. **(a)** Schematic of the GM1 ganglioside synthesis pathway. **(b)** GM3 synthase (*St3gal5*). **(c)** GM2/GD2 synthase (*B4galnt1*). **(d)** GM1 synthase (*B3galt4*). **(e)** β-galactosidase (*Glb1*). For panels (**b-e**), gene expression was analyzed by strain (left) and sex (right). mRNA levels were normalized to *Rpl13a* and are presented as relative expression (mean ± s.e.m). Statistical significance was determined using unpaired two-tailed *Student's t*-test, or two-way ANOVA, as appropriate. * $p \leq 0.05$, ** $p \leq 0.01$, *** $p \leq 0.001$, **** $p \leq 0.0001$. Panel (**a**) was created with BioRender.com.
