## Supplemental Figure 3 for "Cholera toxin remodels intestinal immunity and suppresses systemic metabolism"

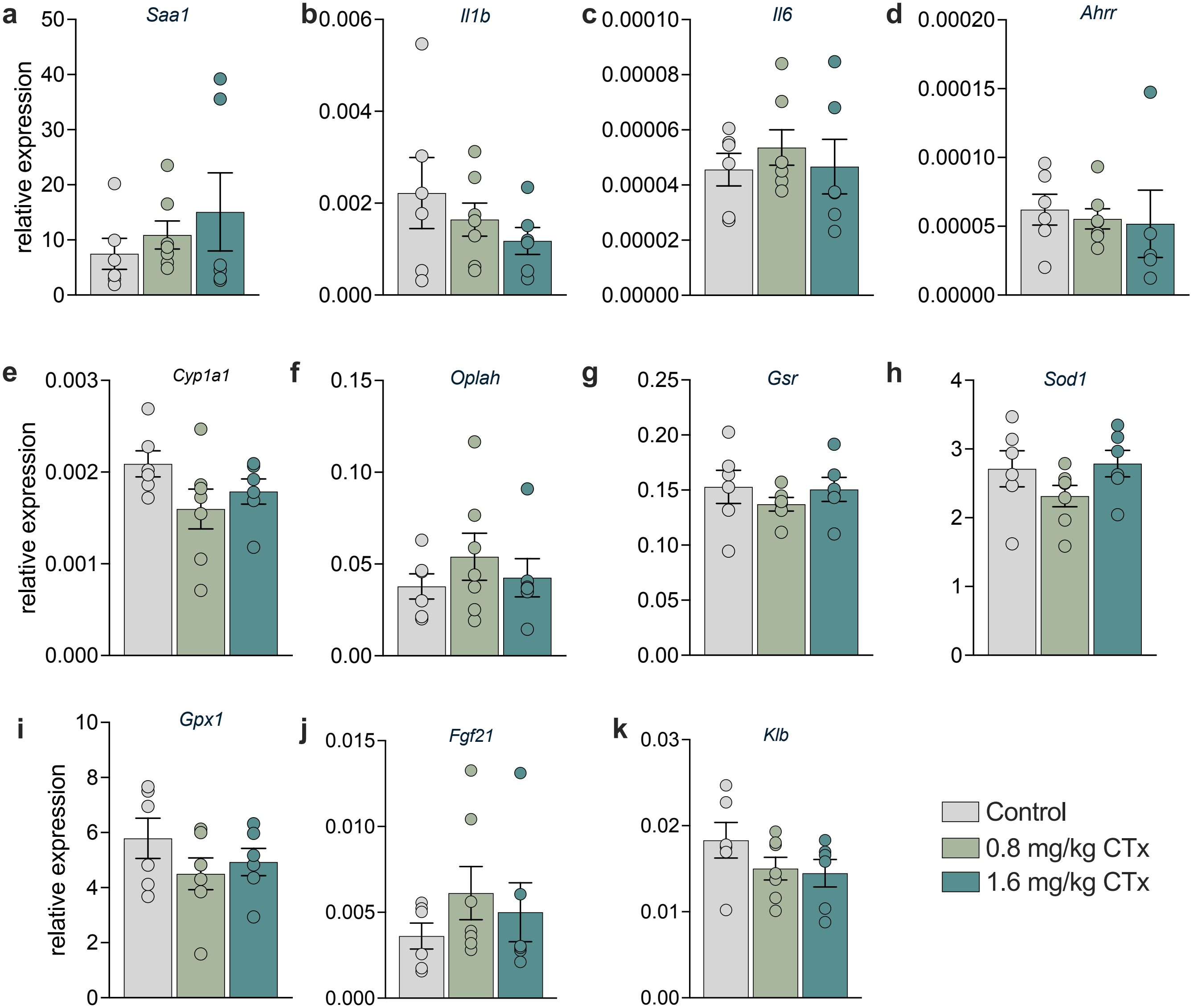


**Supplementary Figure 2 | Hepatic expression of genes not significantly altered following oral CTx exposure.** Livers from C57BL/6 male mice were collected 6 h after a single oral exposure to cholera toxin (CTx) and analyzed by quantitative real-time PCR (n= 6-7 mice per group). Expression levels of the following genes were assessed: **(a)** serum amyloid A (*Saa1*), **(b)** interleukin 1 beta (*Il1b*), **(c)** interleukin 6 (*Il6*), **(d)** aryl hydrocarbon receptor repressor (*Ahrr),* **(e)** cytochrome P450 1A (*Cyp1a1*), **(f)** 5-oxoprolinase (*Oplah*), **(g)** glutathione-disulfide reductase (*Gsr*), **(h)** superoxide dismutase 1 (*Sod1*), **(i)** glutathione peroxidase 1 (*Gpx1*), **(j)** fibroblast growth factor 21 (*Fgf21*), and **(k)** beta-klotho (*Klb*). mRNA levels were normalized to the housekeeping gene *Rpl13a,* and presented as relative expression (mean ± s.e.m). Statistical significance was assessed using unpaired two-tailed *Student's t*-test comparing control and CTx-treated groups. Each panel represents pooled data from three independent experiments. No significant differences were observed between groups.
