## Supplemental Figure 4 for "Cholera toxin remodels intestinal immunity and suppresses systemic metabolism"

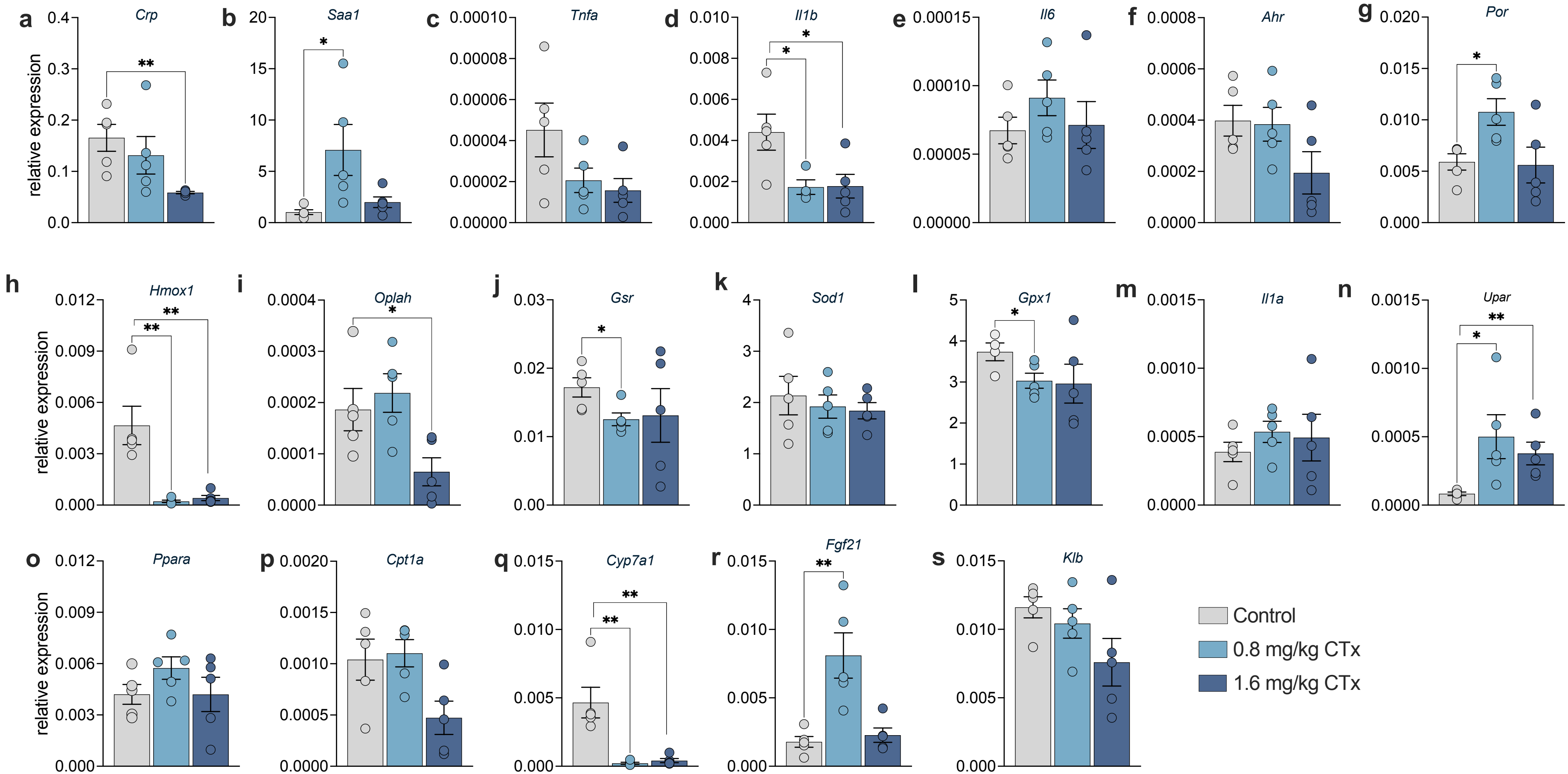


**Supplementary Figure 3 | Hepatic expression of detoxification, oxidative stress and inflammatory mediators following oral CTx exposure in C57BL/6 female mice**. Livers from C57BL/6 female mice were collected 6 h after a single oral exposure to cholera toxin (CTx) and analyzed by quantitative real-time PCR (n= 5 mice per group). Expression levels of the following genes were assessed: **(a)** C-reactive protein (*Crp*), **(b)** serum amyloid A (*Saa1*), **(c)** tumor necrosis factor α (*Tnfa*), **(d)** interleukin-1β (*Il1b*), **(e)** interleukin-6 (*Il6*), **(f)** aryl hydrocarbon receptor (*Ahr*), **(g)** cytochrome P450 oxidoreductase (*Por*), **(h)** heme oxygenase 1 (*Hmox1*), **(i)** 5-oxoprolinase (*Oplah*), **(j)** glutathione-disulfide reductase (*Gsr*), **(k)** superoxide dismutase 1 (*Sod1*), **(l)** glutathione peroxidase 1 (*Gpx1*), **(m)** interleukin-1α (*Il1a*), **(n)** urokinase plasminogen activator surface receptor (*Upar*), **(o)** peroxisome proliferator-activated receptor α (*Ppara*), **(p)** carnitine palmitoyltransferase 1A (*Cpt1a*), **(q)** cholesterol 7α-hydroxylase (Cyp7a1), **(r)** fibroblast growth factor 21 (*Fgf21*), and **(s)** beta-klotho (*Klb*). mRNA levels were normalized to the housekeeping gene *Rpl13a,* and are presented as relative expression (mean ± s.e.m). Statistical significance was determined using unpaired two-tailed *Student's t*-test comparing control and CTx-treated groups. Panels (a-h) show statistically significant differences, whereas panels (i-n) show no significant differences between groups. * p ≤ 0.05, ** p ≤ 0.01. Each panel represents pooled data from two independent experiments.
