## Supplemental Figure 5 for "Cholera toxin remodels intestinal immunity and suppresses systemic metabolism"

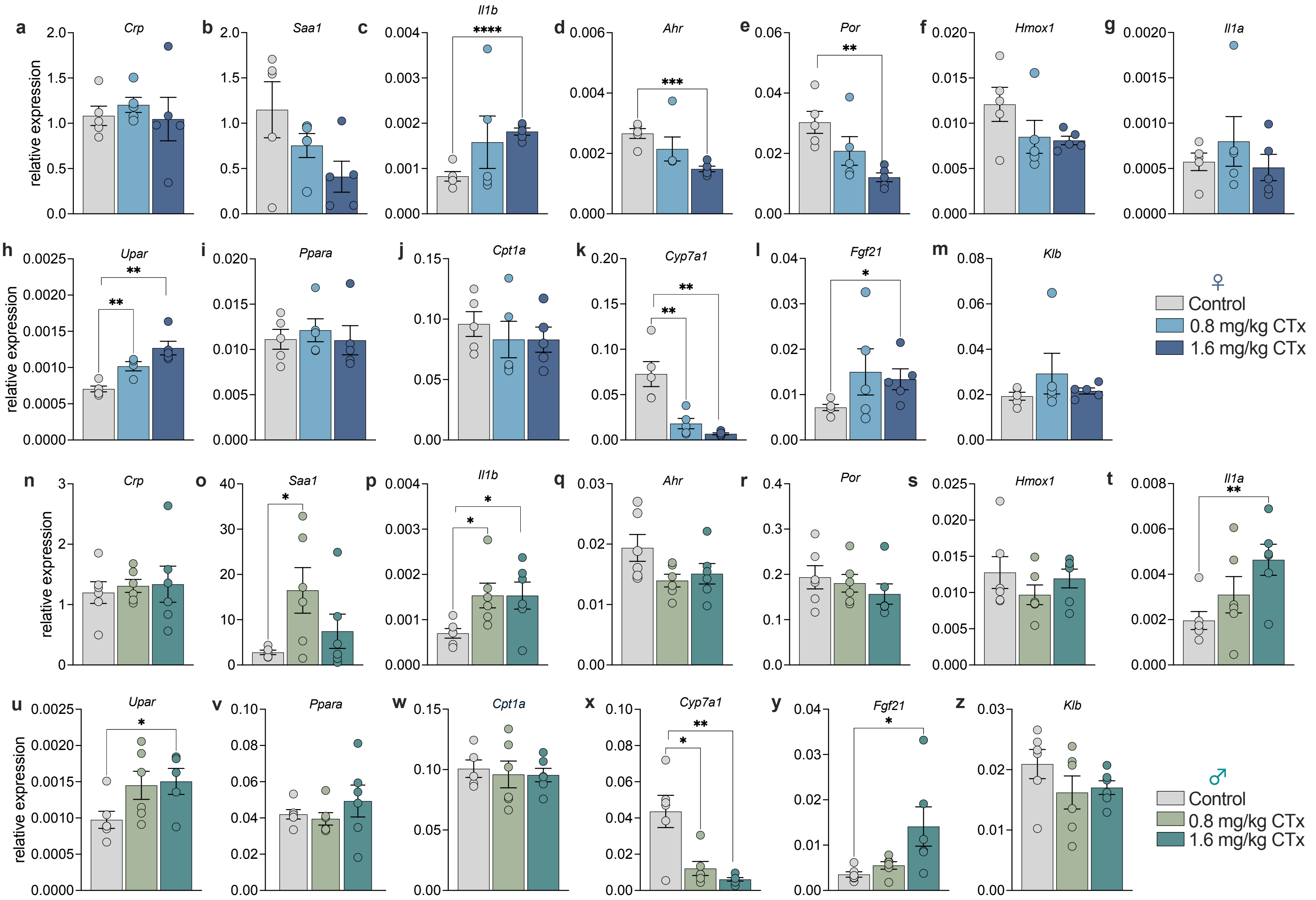


**Supplementary Figure 4 | Hepatic gene expression responses following oral CTx exposure in BALB/c mice.** Livers from BALB/c female (blue bars) and male (green bars) mice were collected 6 h after a single oral exposure to cholera toxin (CTx) and analyzed by quantitative real-time PCR (n= 5-6 mice per group). Expression levels of the following genes were assessed: **(a,n)** C-reactive protein (*Crp*), **(b,o)** serum amyloid A (*Saa1*), **(c,p)** interleukin-1β (*Il1b*)**, (d,q)** aryl hydrocarbon receptor (*Ahr*), **(e,r)** cytochrome P450 oxidoreductase (*Por*), **(f,s)** heme oxygenase 1 (*Hmox1*), **(g,t)** interleukin-1 (*Il1a*), **(h,u)** urokinase plasminogen activator surface receptor (*Upar*), **(i,v)** peroxisome proliferator-activated receptor α (*Ppara*), **(j,w)** carnitine palmitoyltransferase 1A (*Cpt1a*), **(k,x)** cholesterol 7α-hydroxylase (Cyp7a1), **(l, y)** fibroblast growth factor 21 (*Fgf21*), and **(m,z)** beta-klotho (*Klb*). mRNA levels were normalized to the housekeeping gene *Rpl13a* and are presented as relative expression (mean ± s.e.m). Statistical significance was determined using unpaired two-tailed *Student's t*-test comparing control and CTx-treated groups. * $p \le 0.05$, ** $p \le 0.01$, *** $p \le 0.001$, **** $p \le 0.0001$. Each panel represents pooled data from least two independent experiments. Panels (**a-m**) correspond to female mice and panels (**n-z**) to male mice.
