## Supplemental Figure 6 for "Cholera toxin remodels intestinal immunity and suppresses systemic metabolism"

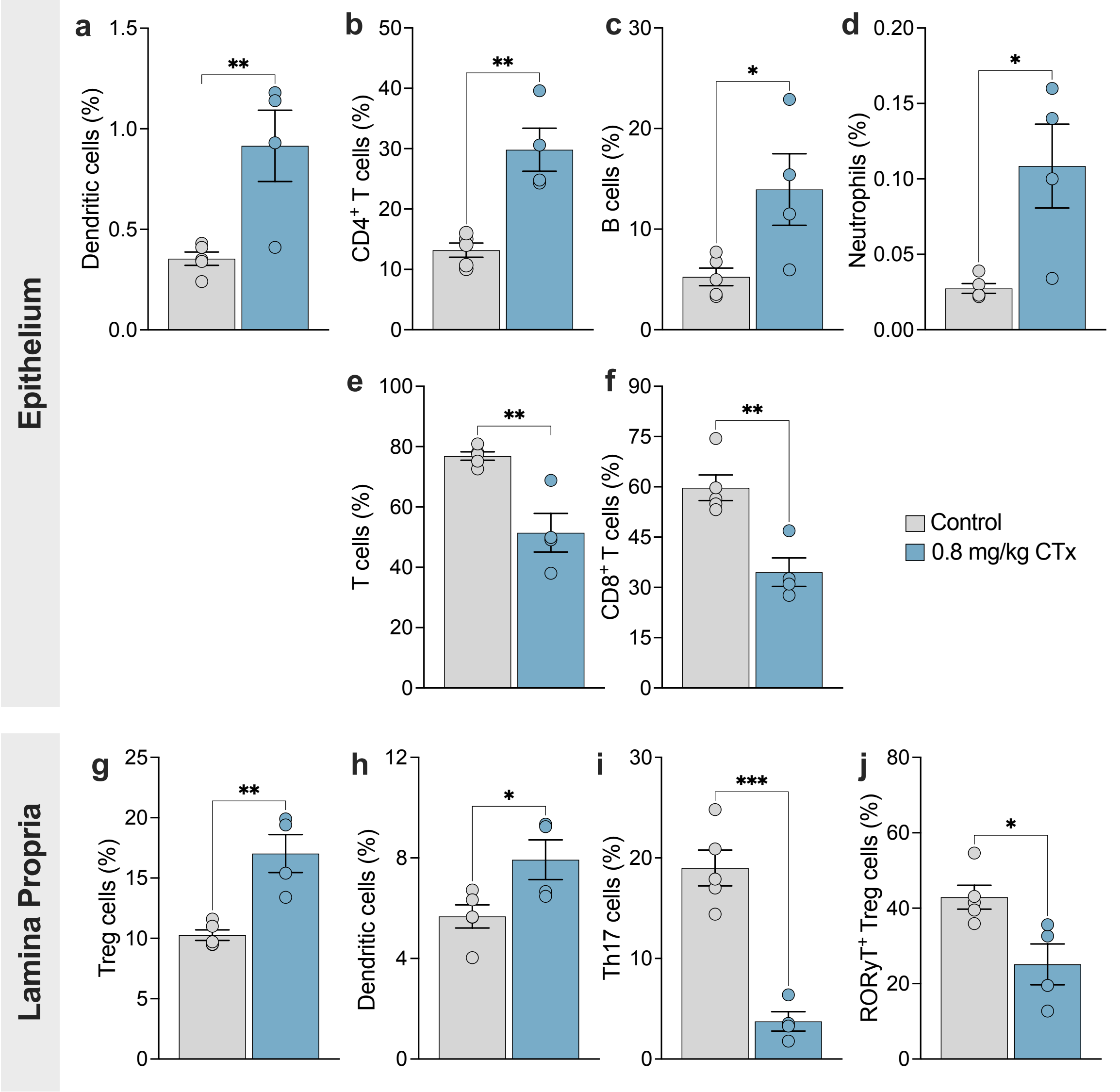


**Supplementary Figure 5 | CTx remodels immune cell composition in the small intestine of C57BL/6 female mice.** Small intestines from C57BL/6 female were collected 24 h after a single oral exposure to cholera toxin (CTx), enzymatically dissociated, and analyzed by full-spectrum flow cytometry (n= 4-5 mice per group). In the intestinal epithelium, CTx changed the frequency of **(a)** dendritic cells ($CD11c^+$ $MHC\text{-}II^+$), **(b)** $CD4^+$ T cells ($CD90.1^+$ $TCR\text{-}\beta^+$ $CD4^+$), **(c)** B cells ($TCR\text{-}\beta^-$ $CD19^+$), **(d)** neutrophils ($CD11b^+$ $Ly6G^+$), **(e)** T cells ($CD90.1^+$ $TCR\text{-}\beta^+$), and **(f)** $CD8^+$ T cells ($CD90.1^+$ $TCR\text{-}\beta^+$ $CD8^+$). In the lamina propria, CTx altered the frequency of **(g)** regulatory T cells (Treg) ($CD90.1^+$ $TCR\text{-}\beta^+$ $CD4^+$ $Foxp3^+$), **(h)** dendritic cells ($CD11c^+$ $MHC\text{-}II^+$), **(i)** Th17 cells ($CD90.1^+$ $TCR\text{-}\beta^+$ $CD4^+$ $CD44^+$ $ROR\gamma T^+$), and **(j)** $ROR\gamma T^+$ Treg cells ($CD90.1^+$ $TCR\text{-}\beta^+$ $CD4^+$ $Foxp3^+$ $ROR\gamma T^+$). All populations were gated on $CD45^+$ live cells. Data are presented as mean ± s.e.m. Statistical significance was determined using unpaired two-tailed *Student's t*-test. * $p \leq 0.05$, ** $p \leq 0.01$, *** $p \leq 0.001$. Data are pooled from two independent experiments.
