## Supplemental Figure 7 for "Cholera toxin remodels intestinal immunity and suppresses systemic metabolism"

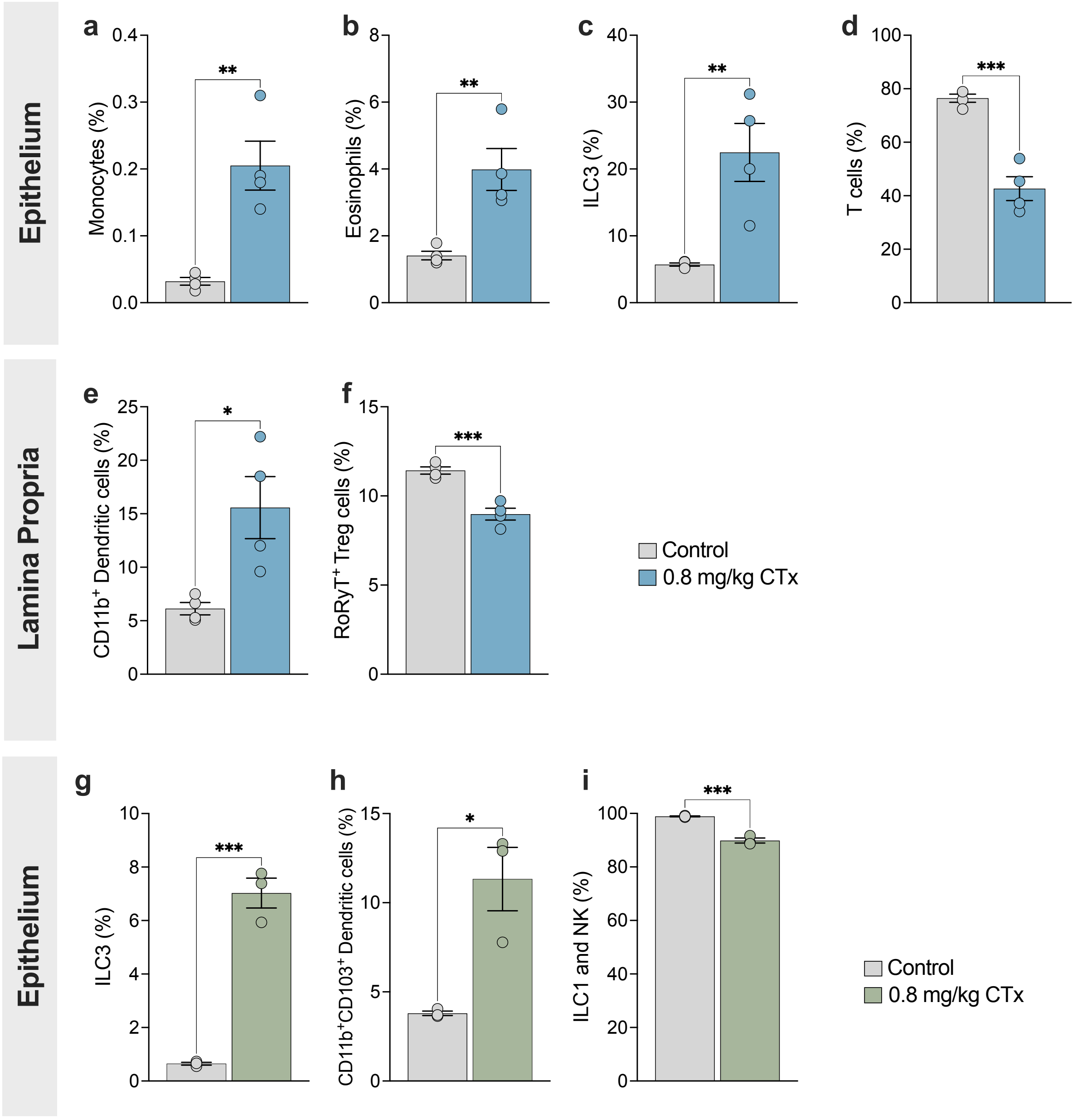


**Supplementary Figure 6 | CTx induces modest changes in immune cell frequencies in the small intestine of mice.** Small intestines from BALB/c female and male mice were collected 24 h after a single oral exposure to cholera toxin (CTx), enzymatically dissociated, and analyzed by full-spectrum flow cytometry. In females, CTx altered the frequency of **(a)** monocytes ($CD11b^-$ $F4/80^-$ $Ly6C^+$), **(b)** eosinophils ($CD11b^+$ $SiglecF^+$), **(c)** ILC3 ($CD90.1^+$ TCR-$\beta^-$ TCR-$\gamma\delta^-$ $RORγT^+$), and **(d)** T cells ($CD90.1^+$ TCR-$\beta^+$). In the lamina propria, CTx increased **(e)** $CD11b^+$ dendritic cells ($CD11c^+$ MHC-$II^+$ $CD11b^+$), and decreased **(f)** $ROR\gamma T^+$ regulatory T cells (Treg) ($CD90.1^+$ TCR-$\beta^+$ $CD4^+$ $Foxp3^+$ $RORγT^+$) (n= 4 mice per group). In males, CTx increased the frequency of **(g)** ILC3 ($CD90.1^+$ TCR-$\beta^-$ TCR-$\gamma\delta^-$ $RORγT^+$), **(h)** $CD11b^+$ $CD103^+$ dendritic cells ($CD11c^+$ MHC-$II^+$ $CD103^+$ $CD11b^+$), and **(i)** ILC1 and NK ($CD90.1^+$ TCR-$\beta^-$ TCR-$\gamma\delta^-$ $GATA3^-$ $RORγT^-$) (n= 3 mice per group). All cell populations were gated on $CD45^+$ live cells. Data are presented as mean ± s.e.m. Statistical significance was determined using unpaired two-tailed *Student's t*-test comparing control and CTx-treated groups. * $p \leq 0.05$, ** $p \leq 0.01$, *** $p \leq 0.001$. Data are pooled from two independent experiments.
